# Cross-linked volumetric DNA microscopy for dense molecular-network phenotyping in intact tissue

**DOI:** 10.64898/2026.06.01.729154

**Authors:** Nianchao Qian, Reem Yasser, Mingrui Yu, Han Chang, Joshua A. Weinstein

## Abstract

Resolving cellular phenotypes in full tissue context requires methods that can retain those cells’ physical neighborhoods, together with the identities of individual biomolecules, in intact three-dimensional specimens. We introduce cross-linked volumetric DNA microscopy (xVDM), in which unique molecular identifiers are seeded directly into the tissue’s protein matrix and linked by uniquely labeled DNA bridges to create a dense, DNA-encoded proximity network. Cell-scale molecular communities are then reconstructed directly from this network. xVDM produces denser molecular networks and broader transcriptome recovery than when these networks are nucleated by transcripts alone. xVDM maps out genetically annotated three-dimensional networks that map onto cell states and tissue regions in intact zebrafish embryos at 12, 18, and 24 hpf. Antibody-oligonucleotide conjugates extend the same framework to protein targets in human tonsil. xVDM provides a route to three-dimensional molecular phenotyping in intact specimens using only standard bench reagents and a DNA sequencer.

## Introduction

Multicellular biology unfolds in three dimensions through cellular proliferation, migration, differentiation, and other phenotypic change that both influence and are influenced by the same breadth of change occurring to their neighbors. Connecting these cells’ gene expression to the emergent physiology of a tissue microenvironment therefore requires methods that can assay identities of both unique cellular phenotypes and unique cellular neighborhoods.

Most spatial-omics technologies approach this problem by capturing or detecting molecules at specific coordinates in Cartesian space. Platforms that generate readouts from either optics or array-capture have made this strategy increasingly powerful at fine resolution ^1;2;3;4;5^. Three-dimensional optical transcriptomic methods have meanwhile extended this direct visualization into thicker tissues ^6;7^. However, volumetric image-capture and cell segmentation remain challenging in dense specimens, particularly in situations where cell boundaries must be inferred from sparse molecular signals ^8^. Physical sectioning extends 2D array-based capture to 3D by imposing a third coordinate in addition to the two already imposed. Optical readouts, meanwhile, broadly preclude untargeted sequencing of single-nucleotide variation, require specialized rigs, and remain subject to the physics of light scattering and collection.

A distinct strategy is to measure molecular connectivity, instead of Cartesian position, across the pre-existing scaffold of biomolecules in an intact specimen. DNA microscopy introduced this idea by encoding local molecular neighborhoods into networks of DNA sequence tags whose pairwise connectivity, once read out by sequencing, could be decoded computationally to recover biologically endogenous ^9;10^ or synthetically patterned geometry ^11;12;13^. Volumetric DNA microscopy (VDM) extended this framework (Fig. 1a–d) to three-dimensional, whole-transcriptome measurements in intact organisms ^14;15^, while seeding molecular tags at transcripts using random primers. Molecular networks form a more “native” language for cell segmentation and tissue biology than pixels ^9;16^. Realizing this potential in tissue, however, requires a molecular network dense enough to survey the specimen’s shape, and not only its most transcript-rich regions. VDM is limited in this respect, because it depends on transcript anchors.

**Figure 1.**
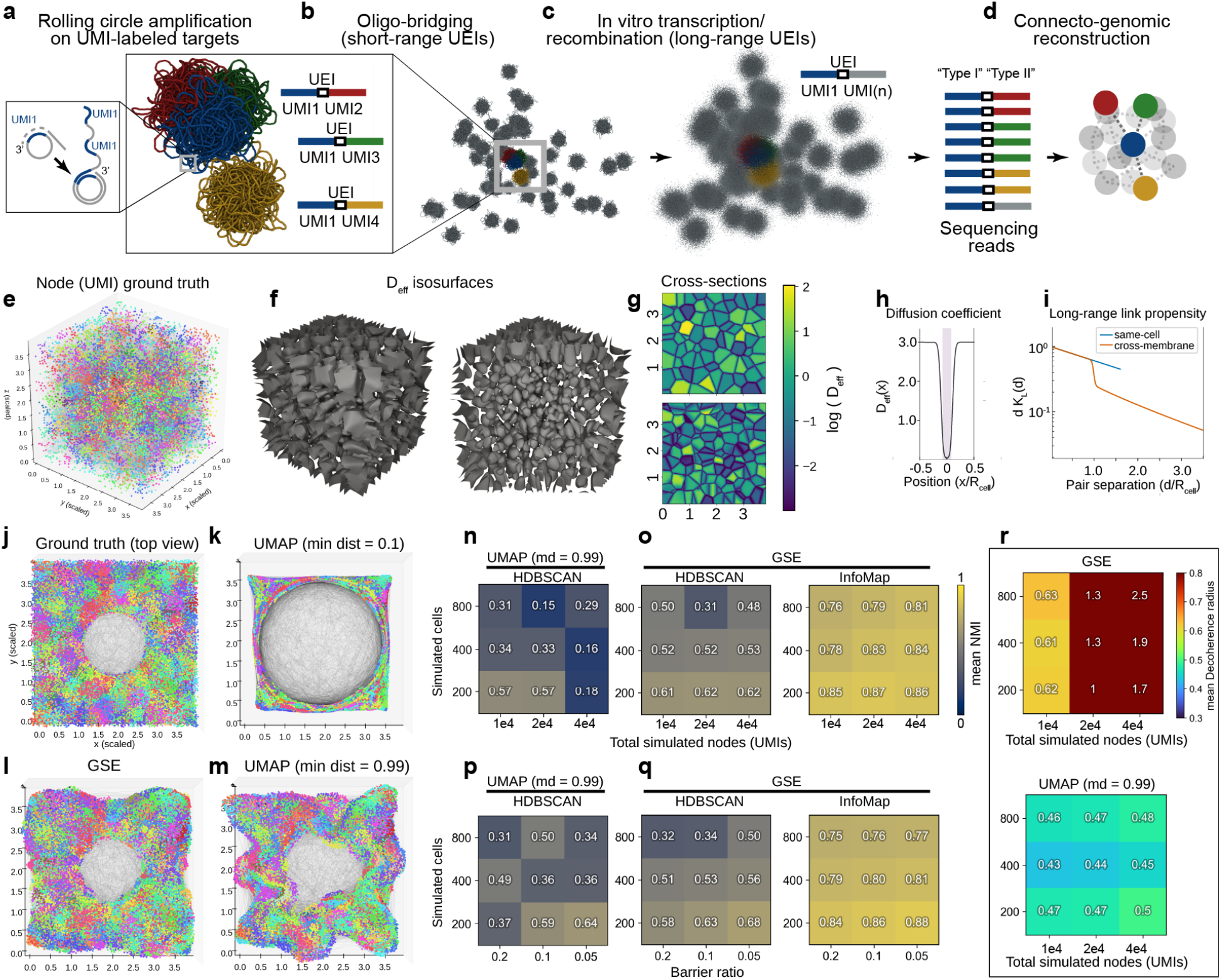
Conceptual framework and simulation benchmark for molecular-network-based cell recovery. **a–d**, Volumetric DNA microscopy concept: rolling-circle amplification on circularized UMI templates creates local DNA nanoballs, bridging oligonucleotides record short-range proximity events, in situ transcription and recombination extend the interaction range, and sequencing of proximity records together with linked molecular identity yields a gene-annotated molecular network. **e**, Ground-truth molecular positions in a Voronoi compartment model. **f**, Representative isosurfaces of the simulated effective diffusion field. **g**, Cross-sections through the same simulated compartments. **h**,**i**, Illustration of membrane-limited diffusion-transport. Two pairs can have the same Euclidean separation but different diffusion-transport history: a path that crosses a membrane contains a narrow low-diffusivity segment, which lowers the path-averaged diffusivity and shortens the effective interaction range. This suppresses long-range linking across compartment boundaries. **j**, Ground-truth simulated node positions, colored by Voronoi cell identity. **k–m**, Graph-based UMAP and GSE embeddings after Procrustes alignment to the simulated positions and colored by the same true labels: UMAP with min_dist=0.1 (**k**), GSE (**l**), and UMAP with min_dist=0.99 (**m**). **n–q**, Mean normalized mutual information to the true Voronoi labels (three replicates), for a joint node- and cell-density sweep (**n**,**o**) and a joint cell-density and membrane-ratio sweep (**p**,**q**). UMAP is scored with HDBSCAN; GSE with both HDBSCAN and Infomap. **r**, Decoherence radius across neighborhood length scales from the node- and cell-density sweep; larger values indicate neighborhoods that stay coherent over longer distances.

Here, we introduce cross-linked volumetric DNA microscopy (xVDM), which overcomes this barrier. xVDM distributes its unique molecular identifiers (UMIs) throughout a specimen’s protein scaffold. It does this by cross-linking short rolling-circle amplification (RCA) primers, via click chemistry, to the tissue matrix independently of where genes are expressed. At each cross-linked primer, RCA copies a circular DNA tag into a localized nanoball carrying many copies of one unique molecular identifier. Neighboring nanoballs are linked to one another by short DNA bridges, the unique event identifiers (UEIs), and to copies of nearby transcripts. Sequencing then yields two coupled readouts from a single specimen: cDNA inserts encode molecular identity (and short-range adjacency, when multiple cDNA inserts link to a single UMI) and UEI bridges encode which molecular nodes were adjacent to one another. Together these readouts form a dense, gene-annotated graph of molecular proximity.

We use zebrafish embryos as a whole-organism test case because they are experimentally tractable as intact specimens, have a well-defined developmental staging system, and are supported by matched-stage dissociated single-cell and spatial transcriptomic references. The 12, 18, and 24 hours post-fertilization (hpf) time points span early organogenesis and somitogenesis while remaining small enough for whole-mount processing, allowing xVDM to be evaluated against both cell-state atlases and coordinate-indexed spatial fields. These analyses focus on what xVDM directly measures, and evaluate the manner and degree to which xVDM supports dense phenotyping of cell-scale structure in intact three-dimensional specimens.

## Results

We use the following vocabulary throughout. A *molecular node* is a UMI-tagged RCA nanoball; a *UEI* ^9;14;15^ is a sequenced proximity record connecting two such nodes; and a *hub* is a molecular node carrying one or more cDNA sub-consensuses. For inference, a *diffusion-transport matrix* is the locally adapted graph used for smoothing and community detection. A *connectivity-derived cell* is an operational unit, a cell-scale molecular community inferred from UEI connectivity and diffusion-transport structure rather than a membrane-resolved anatomical segmentation. Its recoverable boundary depends on graph support: UMI density, UEI yield, molecular contrast, permeabilization, and sequencing depth. A *gene-pole field* is a spatial expression axis defined by marker genes and used for reference-guided registration.

### A simulation benchmark for cell recovery

Recovering cells from a UEI-graph requires an inference pipeline that can convert it into discrete communities of highly inter-connected molecules. To benchmark alternative inference pipelines on varying molecular network densities and topologies, we built a Voronoi-cell model in which molecular nodes were connected by distance-dependent association rules modulated by membrane barriers (Fig. 1e–i). Cellular boundaries within tissue violate the uniformity and isotropy assumptions that underlie the simple DNA microscopy model of reaction-diffusion physics ^9^, making this a controlled test of whether graph structure can still support cell-scale recovery.

VDM had previously demonstrated the effectiveness of using a linear operator – smoothed by a proximity kernel – to construct a linear subspace in which the most likely embedding for molecular nodes could be found ^14^. Doing so led to notable improvements in reconstruction over short and long length scales, quantified by decoherence analysis (in which displacement-errors are plotted as a function of difference from randomly chosen reference points), compared to earlier subspace methods ^9^ and UMAP ^17^. This smoothing-and-embed procedure, called Geodesic Spectral Embedding (GSE), used fixed Gaussian-proximities that, though useful, could not be interpreted as reflective of the data itself (in much the same way nearest neighbor graphs writ-large are arbitrary deformations of high-dimensional data ^18^).

To resolve this, and improve the robustness of the proximity kernel within GSE, we replaced the fixed Gaussian kernel with a well-defined objective function whose solution encoded each individual UEI connection as a linear sum of nearest neighbor positions in the initial UEI-matrix eigenvector subspace (*SI: Stabilized geodesic kernel*). This was motivated by recent empirical success in mitigating DNA microscopy noise by down-weighting anomalous UEI edges ^19^. Specifically, for each UMI-node, we fit each UEI-edge displacement as a weighted average of nearest-neighbor displacement vectors, letting each local connection experienced by a UMI adapt toward network neighborhoods most strongly supported by the dataset as a whole. This transport matrix was then used for UEI-matrix smoothing in the same way as before ^14^, followed by maximizing the inferred-embedding likelihood and segmentation.

We benchmarked UMAP ^17^ against GSE using identical inputs and downstream segmentation methods. For UMAP, we included the high-min-dist parameterization recently proposed for DNA microscopy on uniform arrays ^11^. The two segmenters use different information sources: HDBSCAN ^20^ models node density in embedding coordinates, whereas Infomap ^21^ models node diffusion along an inputted transport graph. In our case, GSE’s transport matrix (*SI: Stabilized geodesic kernel*) compresses information about both local neighborhoods within the embedding and observed UEI connectivity, so Infomap can in principle access information about an xVDM dataset’s structure that density-based segmentation cannot.

Fig. 1j shows the ground-truth simulated positions, and the graph-based UMAP and GSE embeddings in Fig. 1k–m are colored by the same true Voronoi labels rather than by cluster calls. They show that global shape recovery and the recovery of cellular identities are separable objectives. UMAP can preserve the outer outline while broadening local neighborhoods in a way that degrades density-based segmentation (Fig. 1n,p). GSE sharpens those same neighborhoods into denser, more isolated molecular nodes, so the identical HDBSCAN procedure performs better after GSE than after UMAP (Fig. 1o,q). Infomap’s higher score under matched embeddings (Fig. 1o,q) and in reconstructing simulated mixed populations of cells (Fig. S4) reflects its use of both the embedding and the graph displacements that shaped the learned diffusion-transport matrix. This motivates our choice to call cells in real xVDM data using Infomap on the diffusion-transport matrix.

A decoherence analysis ^14^ – which measures how reliably local neighborhoods are recovered as the length scale changes – showed that GSE’s advantage over UMAP grew with network size (Fig. 1r). The simulation therefore makes the methodological claim that given a sufficiently dense network from whole-embryo xVDM, GSE embedding followed by Infomap on the diffusion-transport matrix can recover cell-scale modularity from connectivity alone. In tissue, the corresponding objects are connectivity-derived molecular communities, and their anatomical correspondence is evaluated through the independent molecular and spatial benchmarks that follow.

### Matrix seeding densifies the network

We next isolated the chemistry question: whether protein-matrix seeding supplies the graph density and local molecular sampling needed for the computational pipeline to operate in real tissue. We applied xVDM to whole zebrafish embryos at 12, 18, and 24 hpf, cross-linking amine-modified RCA primers to the tissue protein matrix to seed UMIs independently of local gene expression. Physically, this chemistry converts the fixed tissue matrix into a dense set of localized molecular sampling sites. Each cross-linked RCA primer seeds a DNA nanoball that remains localized near its anchoring point and carries many copies of the same UMI. Bridging oligonucleotides then record short-range contacts among neighboring nanoballs and link nearby cDNA products to those local UMIs. The result is a sequencing-readable network in which matrix-anchored molecular nodes carry both proximity information and local molecular identity.

Fluorescent-dUTP incorporation during RCA confirmed pronounced nanoball signal from circularized UMI templates in whole-mount 24 hpf embryos but not from linear UMI controls (Fig. 2a). Improved yield in mammalian tissue was achieved by coupling oligonucleotides through activated thiol groups (Fig. 2b). Applying the chemistry end-to-end (Fig. 2c–k) generated UEI-amplicon and cDNA-amplicon sequencing libraries. UEIs and UMIs were analyzed separately as previously described ^14^, and the cDNA reads belonging to each UMI in the cDNA library were grouped into sub-consensuses (Fig. 2k). Each sample’s UMIs belonging to reads in its cDNA library were matched to UMIs in its UEI library. Although retention of genes through this filtering process was broadly uniform, epidermal marker genes were found to most effectively survive the data pruning process (Fig. S5).

**Figure 2.**
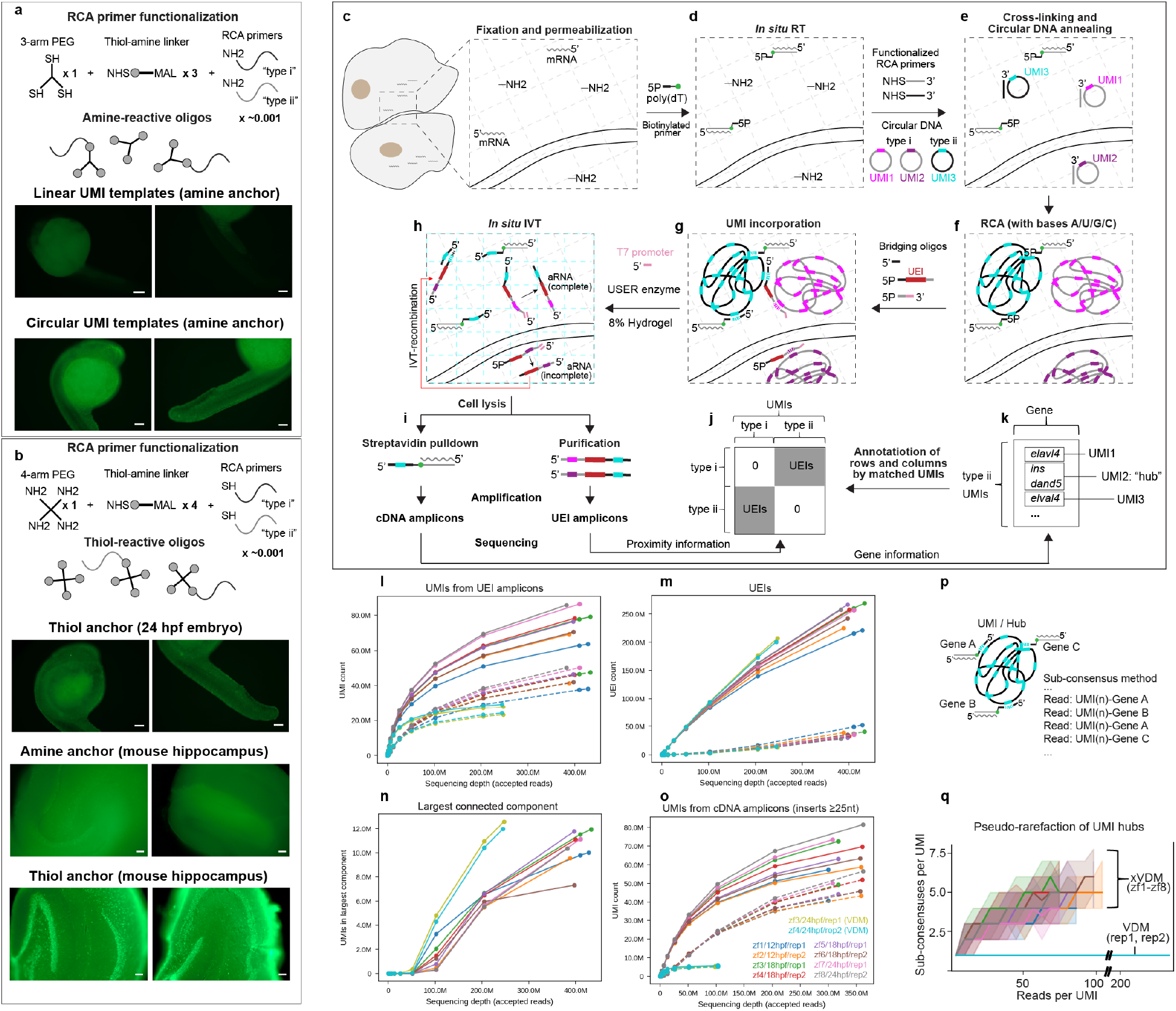
xVDM densifies the UMI network. Samples are fixed and permeabilized to preserve the spatial localization of mRNA and proteins while rendering them accessible for subsequent *in situ* reactions. (**a**) RCA primers bearing 5’ amine (NH_2_) are covalently cross-linked to free NH_2_ groups within the sample via thiolated multi-arm PEG and subsequently annealed to either circular or linear RCA/UMI templates, with signal read out via fluorescent dUTP incorporation. (**b**) Alternatively, cross-linking is performed using thiolated (SH) RCA primers and amine-functionalized PEG. In the end-to-end xVDM reaction, mRNA is reverse transcribed using a poly(dT) primer with an internal biotin modification (solid green circle) and a 5’ phosphate (5’P), and UMIs are locally amplified by cross-linked RCA into DNA nanoballs (**c–f**). **g**, Bridging oligonucleotides hybridize to the nanoballs, incorporating UMIs into both UEIs (unique event identifiers) and cDNA. **h**, UEI products are further amplified *in situ* through hydrogel-embedded *in vitro* transcription (IVT). At low frequency, amplified RNA (aRNA) derived from intermediate UEI products undergoes templated recombination via unanchored diffusion (red arrow). **i**, Biotinylated cDNAs are recovered by streptavidin pulldown to capture gene identity (**k**), while IVT-derived UEI amplicons (**j**) encode molecular proximities. Rarefaction shows boosted yield over VDM (**l–o**). By allowing multiple cDNA inserts to anneal to the same UMI simultaneously, each UMI constitutes a “hub” for gene-insert detection (**p**). Sub-consensus counts accumulate with increasing read depth (**q**).

At matched sequencing depth, xVDM recovered substantially more molecular nodes, more cDNA bearing hubs, and more genes than transcript anchored VDM at 24 hpf (Fig. 2l–o, Table S2). Rarefying UEI amplicons to 204.8 million accepted reads, xVDM yielded 2.5× more UEI associated molecular nodes than VDM (69.0 versus 27.8 million). Rarefying cDNA amplicons to 51.2 million accepted reads, xVDM yielded 6.2× more cDNA bearing molecular hubs (32.6 versus 5.3 million) and 1.4× more genes at ≥ 10 weighted support (21,592 versus 15,738) (Fig. 2l–o, Fig. S3, Table S3). These matched depth gains establish the central chemistry claim independently: matrix seeding increases both the number of molecular nodes that enter the proximity graph and the number of graph nodes that carry local transcript identity.

### cDNA hubs capture local molecular context

Because cDNA hubs are molecular nodes within the sample at which one or more local cDNA inserts colocalize, they provide a direct readout of short range molecular adjacency before reconstruction (Fig. 2p). For each UMI, we counted the number of cDNA sub consensuses, each gated at ≥ 2 supporting reads, against the reads used to generate them (Fig. 2q). Processed through the same pipeline, the older VDM data yielded only one gene per tag even at high read depth, confirming that the pipeline does not artificially fuse unrelated genes. In xVDM, hubs accumulated multiple sub consensuses as read depth increased, consistent with protein matrix seeded UMIs acting as local samplers of nearby cDNA molecules.

We used cDNA hub composition as a control on local molecular context. Within each embryo, hubs containing cytosolic rRNA and no mitochondrial rRNA were matched to hubs containing mitochondrial rRNA and no cytosolic rRNA by UMI type and detected-feature-count bin. Across the eight embryos, inverse-variance fixed-effect meta-analysis of embryo-level log odds ratios, followed by Benjamini–Hochberg adjustment over tested protein-coding genes, identified 38 genes enriched in cytosolic-rRNA hubs and 19 genes enriched in mitochondrial-rRNA hubs at *q* < 0.05. Mitochondrial-rRNA hubs were enriched for mitochondrial transcripts. Cytosolic-rRNA hubs favored lipid and apolipoprotein genes, including *apoa1a, afp4, apoa1b, tfa*, and *fabp1b*.*1* (Fig. 3a). The per-embryo profiles and the 24 hpf minus 12 hpf comparison show how this compartment effect varies across specimens and developmental stages (Fig. S6a,b).

**Figure 3.**
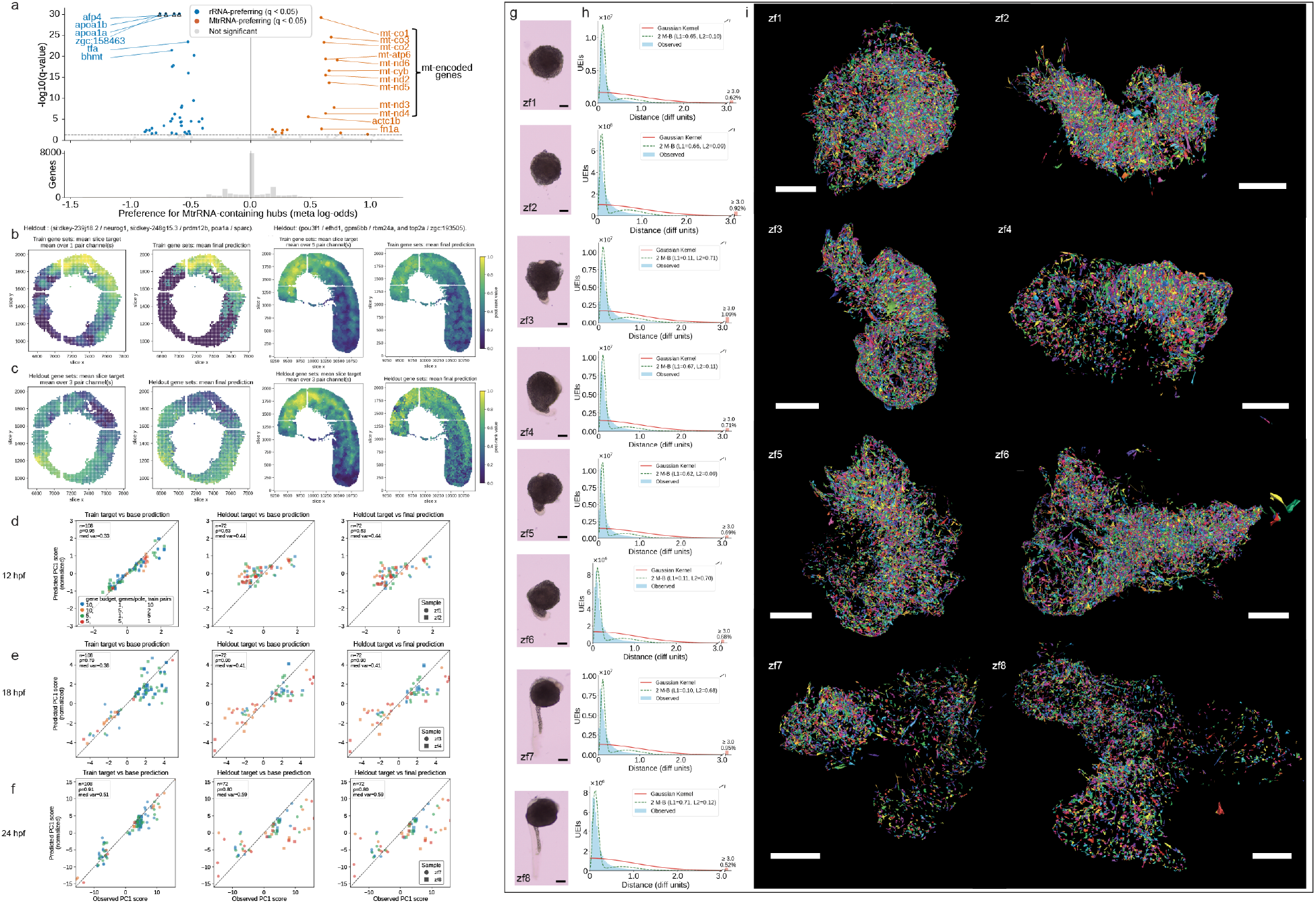
xVDM preserves molecular and spatial information over multiple length scales. (**a**) cDNA-hub compartment analysis. Hubs containing cytosolic rRNA and no mitochondrial rRNA were matched within each embryo to hubs containing mitochondrial rRNA and no cytosolic rRNA by UMI type and detected feature count. Points are protein-coding genes. The x-axis gives the inverse-variance fixed-effect log odds ratio for mitochondrial-rRNA hubs relative to cytosolic-rRNA hubs. Positive values indicate mitochondrial-rRNA association, and negative values indicate cytosolic-rRNA association. The y-axis shows − log_10_(*q*_display_), where *q*_display_ = max(*q*, 10^−30^) is used only for plotting and *q* is the Benjamini–Hochberg adjusted value over tested protein-coding genes. Colored genes pass the unclipped fixed-effect *q <* 0.05. **(b**,**c)** Example reference-guided registration fields at 12 hpf and 18 hpf, each shown for one representative training setting. Each pair shows the Stereo-seq target field (a rank-transformed map of where one gene group is over-represented relative to an opposing group) and the xVDM prediction after registration. Fields in **(b)** were used to fit the assignment; fields in **(c)** were held out and used only for evaluation. **(d–f)** PC1 correlograms summarizing registration at **(d)** 12 hpf, **(e)** 18 hpf, and **(f)** 24 hpf (PC1 of the registration benchmark; full sweeps in Fig. S8). Within each run the target fields define the PC1 axis; axes are target vs. predicted PC1 score, and the diagonal marks perfect agreement. **(g)** Bright-field images of specimens zf1–zf8. **(h)** Connection-spread functions: UEI-weighted distributions of GSE distances between molecular nodes joined by retained UEI edges. A one-component Gaussian-dispersion reference is shown with a fitted two-component Maxwell–Boltzmann mixture (Methods). The short component reports local molecular spread and the broader reports graph-spanning connectivity. Percentages at right give the UEI mass at distances ≥ 3 diffusion units, outside the plotted range. **(i)** xVDM surface renderings randomly colored by cell identity. Bright-field scale bars, 200 µm; xVDM scale bars, 25 diffusion units ( ∼ 10 µm per diffusion unit, ^14^).

The same hub sets also showed patterns of gene-modules across developmental groups. In hubs with cytosolic rRNA, lipid genes and protein synthesis genes were detected together more often than genes drawn from the two groups. In hubs with mitochondrial rRNA, nuclear translation genes showed the strongest same group signal in most groups, while mitochondrial genes were closer to the between group comparison (Fig. S6c).

### A connectivity-based resolution measure

We next asked how rapidly molecular proximity falls off with inferred distance in each xVDM reconstruction. We adopt the term *connection-spread function* (CSF) for the UEI-weighted distribution of GSE distances between molecular nodes joined by retained UEI edges. Because xVDM infers geometry from a connectivity graph, the CSF can be understood as a counterpart to an optical point-spread function but at the level of connectivity, reporting how rapidly retained proximity signal decays in the inferred geometry. Under the locally uniform Gaussian-dispersion model previously used in transcript-anchored VDM ^14^, the expected radial form is Maxwell–Boltzmann, ∝ *r*^2^ exp( − *r*^2^/*L*^2^).

Across all eight embryos, the observed CSFs exhibited a sharp local peak together with a broader tail. A one-component Gaussian-dispersion/Maxwell–Boltzmann reference did not capture both regimes, whereas a two-component Maxwell–Boltzmann mixture summarized them with a short component at 0.09–0.12 diffusion units and a broader component at 0.6–0.7 diffusion units (Fig. 3h). Under the empirical scale conversion of ∼10 µm per diffusion unit established for VDM ^14^, the short component corresponds to a micron-scale local proximity kernel and the broader component to longer-range connectivity that keeps the molecular graph contiguous across the specimen.

### Registration to a spatial reference

We then asked whether xVDM cDNA also carries spatial structure that can be matched to an independent, coordinate-indexed reference. To do this, we generalized the Type A/Type B construction ^14^ to many *gene-pole fields*: from a matched-stage Stereo-seq slice, each field reports where one small gene group is locally over-represented relative to an opposing gene group. We used a subset of these fields to map aggregated xVDM cells onto specific sites within the Stereo-seq slice, holding out the remaining fields for evaluation. At 12 hpf and 18 hpf, post-registration xVDM predictions reproduced the slice’s spatial gradients in both training and held-out fields (Fig. 3b,c). Compressing this comparison across embryos onto the leading target-defined axis of spatial variation gives the PC1 correlograms in Fig. 3d–f: held-out fields landed near the diagonal at all three stages, and UEI-matrix refinement did not degrade this transfer. PC1 captured 44.1% of held-out target-field variance at 12 hpf, 43.5% at 18 hpf, and 60.7% at 24 hpf, so it is a partial but informative summary at early stages and a stronger summary once the embryo has elongated along a dominant body axis.

### Expression states map to tissue programs

We next asked whether cell states inferred from xVDM’s molecular communities correspond to anatomically meaningful tissue programs. We aggregated putative cells, implementing a quality-control filter that required each to contain at least 100 zebrafish transcripts that could be uniquely mapped to the genome and 50 distinct genes. In all eight embryos, we normalized and log-transformed the resulting per-cell cDNA count matrix, selected highly variable genes with specimen as the batch covariate, and computed a single principal-component embedding, a 30-nearest-neighbor graph over 30 principal components, and one Leiden clustering ^22^ of the pooled object. The Leiden expression labels, *L*_0_–*L*_15_, are therefore shared across the analysis rather than independently fitted within each specimen. For visualization, these same pooled labels were rendered separately in each embryo (Fig. 4a, left of each pair). After coarsening and registering each reconstruction to a same-stage Stereo-seq slice (Fig. 4a, right of each pair), the expression-labeled regions aligned with stage-appropriate tissue annotations from ^23^: neural rod, adaxial cells, segmental plate/tail bud, somites, and polster at 12 hpf; notochord, neural crest, and erythroid lineage at 18 hpf; and musculature, fast muscle, and dorsal/ventral spinal cord at 24 hpf.

**Figure 4.**
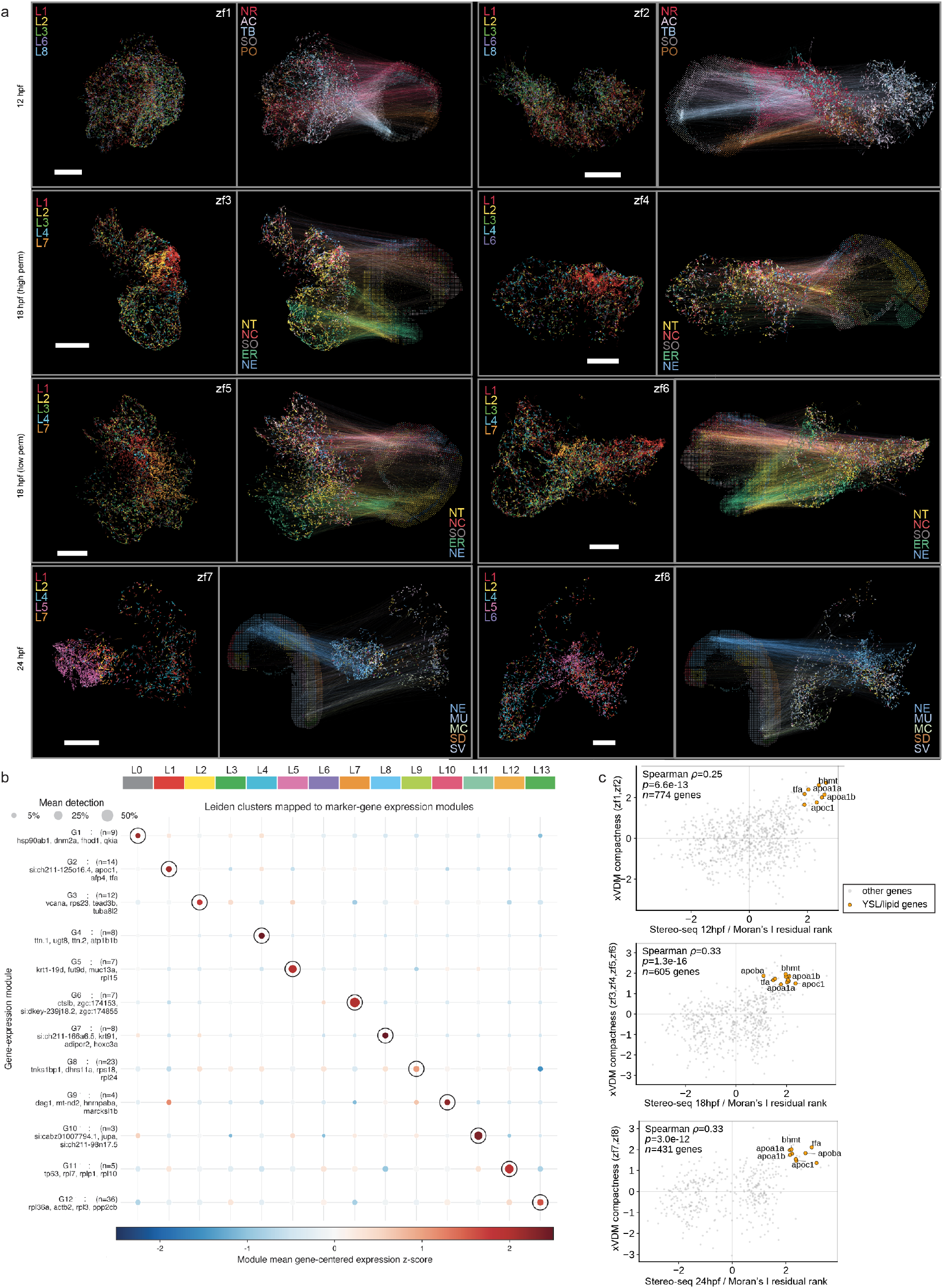
Expression labels, tissue-reference alignment, and gene-level spatial validation of xVDM reconstructions. **(a)** Paired xVDM renderings of each specimen. The left member is colored by the pooled Leiden expression label *L*_*k*_ computed from the per-cell cDNA matrix across all eight embryos; labels are shared across the analysis, although not every label appears in every specimen. The right member shows the same xVDM view alongside a matched-stage Stereo-seq slice. Arcs connect plotted aggregate cells to their expected positions on the registered slice after coarsening, colored by the slice tissue annotation at the supported locations ^23^. Rows are 12 hpf, 18 hpf high-permeabilization, 18 hpf low-permeabilization, and 24 hpf specimens. xVDM scale bars, 25 diffusion units. Tissue abbreviations: NR, neural rod; AC, adaxial cell; TB, segmental plate/tail bud; SO, somite; PO, polster; NT, notochord; NC, neural crest; ER, erythroid lineage; NE, nervous system; MU, musculature/yolk syncytial layer; MC, fast muscle cell; SD, spinal cord, dorsal; SV, spinal cord, ventral. **(b)** Dot plot relating pooled Leiden expression labels (columns) to marker-gene modules (rows). The leftmost text lists representative top genes. Dot color is the within-module mean of gene-centered expression *z*-scores. Dot size is the mean fraction of cells in which module genes are detected. Black circles mark each module’s dominant label. **(c)** Gene-level spatial validation at 12 hpf, 18 hpf, and 24 hpf, shown as three stage-specific subpanels. Each point is one gene. The *x*-axis is the rank-normalized residual of external Stereo-seq Moran’s *I*, where Moran’s *I* measures spatial autocorrelation in the matched-stage reference slice. The *y*-axis is the rank-normalized residual of xVDM gene compactness, where larger compactness means that high-expressing aggregate cells for that gene are closer to one another in inferred three-dimensional xVDM space than expected from the specimen-wide background. Residualization removes external and xVDM mean expression, detection frequency, and expression variance. Insets report the Spearman rank correlation coefficient *ρ*, the two-sided *p* value, and the number of genes *n*.

The marker-module map (Fig. 4b) gives an interpretation at the gene-level of these pooled expression labels. Each column is one pooled label and each row is a marker-gene module selected from balanced cluster-versus-rest expression contrasts. Module G4, containing genes such as *ttn*.*1, ttn*.*2, ugt8*, and *atp1b1b*, reports a muscle-associated program. A separate lipid- and yolk-associated module, enriched for apolipoprotein and yolk-program genes including *apoc1, afp4*, and *tfa*, marks the yolk-syncytial-layer/lipid program, whereas a distinct hatching-gland/protease-like module is dominated by genes such as *ctslb*. The pooled labels in Fig. 4a are therefore compact expression-state labels whose marker modules support tissue-level interpretation.

As a separate check on the *ctslb* marker module in Fig. 4b, we tested cDNA hubs from the two 24 hpf embryos. Hubs carrying *ctslb* were matched within each 24 hpf embryo to control-hubs lacking *ctslb* by UMI type and detected-feature-count bin. Hubs with *ctslb* were more likely to also carry cDNA from *si:dkey-269i1*.*4, zgc:174855, zgc:174153*, and *zgc:158463*. The *hbae3* anchor did not show a clear set of non hemoglobin genes. These results support local cDNA sharing within the *ctslb* marker module, but they do not imply binding or regulation (Fig. S7).

### Gene-level spatial structure is preserved

We next asked whether xVDM recovered spatial organization at the level of individual genes, rather than only at the level of expression labels. For each developmental stage, we computed Moran’s *I*, a spatial autocorrelation statistic, for genes in the matched-stage Stereo-seq reference slice using the reference’s physical two-dimensional coordinates. Independently, in the stage-matched xVDM embryos, we computed a gene compactness score that increases when aggregate cells with high expression of that gene lie closer to one another in inferred three-dimensional xVDM space than expected from the specimen-wide background. Compactness was computed globally within each specimen, not within a metacluster, and gene-level compactness scores were averaged across the stage-matched specimens.

Spatial autocorrelation and compactness can both be affected by expression level, detection frequency, and expression variance. We therefore compared rank-normalized residuals after regressing both the external Stereo-seq and xVDM measurements against these covariates. Genes with high Stereo-seq spatial autocorrelation were also compact in xVDM at the same developmental stages (Fig. 4c). In the 18 hpf subpanel of Fig. 4c, for example, external Moran’s *I* and xVDM compactness were positively correlated (Spearman rank correlation coefficient *ρ* = 0.33, two-sided *p* = 1.3 × 10^−16^, *n* = 605 genes), and YSL/lipid genes including *apoc1, tfa, apoa1a, bhmt, apoa1b*, and *apoba* occupied the high–high quadrant. The inferred xVDM geometry therefore preserves gene-level spatial organization measured independently by Stereo-seq.

### Targeting proteins with xVDM in human tonsil

We next tested the full xVDM workflow in mammalian tissue, in particular whether the TCEP-coupling already optimized in a mammalian hippocampus (Fig. 2b) could be extended to oligonucleotide tagged protein targets in human tonsil cryosections. We sought to use antibody-oligonucleotide conjugates (AOCs) developed for Prox-Seq ^24;25^ that could incorporate a proximity-ligation signal directly into the cDNA libraries that could, in turn, be amplified as part of xVDM’s readout. This would align with concurrent efforts to incorporate protein identities into DNA microscopy readouts ^10;26^, but extend this body of methods to multi-cellular tissue.

In sections from two human tonsil specimens, libraries showed rapid saturation of antibody proximity-ligation UMIs and slower saturation of UEI libraries (Fig. S9). For the cDNA-library hub analysis, we retained antibody-derived inserts whose sequence contained exactly two recognized epitope motifs, assigning each retained insert an epitope-pair label such as CD3/CD3, CD3/IgD, or IgD/IgD. Among hubs carrying exactly two retained inserts, same-epitope co-localization across distinct inserts, including CD3 with CD3 and IgD with IgD, was enriched after false-discovery-rate control in permutation tests that shuffled epitope-pair labels among the two fixed insert slots per hub while preserving the global abundance of each label (Fig. S10a,d). Restricting the label set to homo-epitope pairs, where labels such as CD3/CD3 and IgD/IgD denote inserts carrying two motifs from the same epitope class, gave a concordant result (Fig. S10b,e). These results show that antibody-derived tags preserve local protein context at hubs, in the same embedding-independent sense that cDNA hubs preserved local transcript context in zebrafish embryos (Fig. 3a, Fig. S6).

The UEI-based neighborhood analyses used hubs with exactly one epitope-motif-dimer (i.e., qualifying) sequence insert, denoted *d* = 1, where *d* is the number of qualifying inserts on a hub. In *K*-nearest-neighbor analyses, where *K* is the number of neighboring hubs included in each local neighborhood, CD3/CD3 and IgD/IgD signals separated across neighborhoods. Infomap-cluster maps using *K* = 30 showed CD3-rich and IgD-rich regions in distinct parts of the inferred tonsil maps, consistent with T cell and B cell zones (Fig. S10c,f,g,h). This experiment, though not designed as a complete tonsil atlas, showed xVDM capable of mapping protein organization, via antibody tags, in clinical specimens.

## Discussion

xVDM advances volumetric DNA microscopy by pairing improvements in both chemistry and computational inference. By departing from the requirement that each UMI must be seeded at transcripts, and instead using protein matrices within fixed specimens as a scaffold, xVDM produces a denser proximity matrix and hubs that can accommodate a multiplicity of gene inserts, improving the detection of genetic diversity and localization across intact samples. xVDM image inference is meanwhile adapted to the resulting larger and more modular molecular graphs that result. Network analysis aggregates these molecular nodes into putative cells, and solving a capacity-constrained registration problem allows these putative cells to be mapped to 2D array-based spatial transcriptomic platforms.

xVDM reconstruction captured biologically meaningful spatial organization at three distinct length scales. At the sub-cellular level, rRNA-hub composition recovered cytosolic-versus-mitochondrial partitioning before any embedding. At the gene level, genes with the highest spatial autocorrelation in a 2D commercial platform (Stereo-seq) were also those with the most compact spatial distributions in xVDM coordinates. At the tissue level, held-out gene-markers, when expressed as spatial fields along the dominant axis of coordinates in 2D slices of zebrafish at the same embryonic age, mapped onto those inferred by xVDM coordinates. Collectively, these results established how the xVDM framework can be a force-multiplier in the reconstruction of tissue-level biology alongside platforms with different sensitivities and biases.

xVDM occupies a distinct point in the spatial-omics space. Compared with array-based capture ^1;4;5^, it provides three-dimensional reconstruction without sectioning. Compared with 3D imaging-based transcriptomics ^7;6^, it removes the requirements for optical access, panel design, and segmentation in dense tissue, at the cost of direct optical visualization, but distinguishing single molecules by downstream “de novo” genic sequencing. xVDM is therefore most useful where the specimen is intact and three-dimensional, where optical access is limited, and where unrestricted transcript capture is preferred over a curated panel.

Resolution in xVDM is most naturally defined in the space formed by the UEI graph. Because geometry is inferred from connectivity, no single Abbe-like number applies. Instead the resolving power on each specimen is empirically determined by a connection-spread function (Fig. 3h). Under the empirical VDM/xVDM scale correspondence of roughly 10 µm per diffusion unit, recapitulated in this paper by side-by-side comparisons of bright-field and xVDM scale bars, the short CSF component corresponds to approximately micron-scale local molecular spread. The broader component reflects the longer-range connectivity that keeps the graph contiguous across the specimen. Both what CSF is achievable and the sharpness of its boundaries depend on UMI density, UEI yield, permeabilization, and sequencing depth.

This connectivity further defines how, in its current generation, xVDM’s putative inferred cells can be interpreted, namely as communities within the inter-molecular network that will mirror cell boundaries only insofar as the UEI-graph’s density allows it. xVDM’s pushing this density upward makes it a critical waypoint in the advance toward single-cell-resolved volumetric DNA microscopy. The impact of this improvement in density can be seen through three distinct components of the data: sub-cellular structure of hubs carrying rRNA (Fig. 3a), the CSF falloff of UEI edges (Fig. 3h), and transfer of held-out gene-pole fields and same-stage tissue annotations after reference-guided registration (Figs. 3b–f, 4a). Improved UEI-yield modeling, more uniform molecular seeding, and joint protein/transcript readout should sharpen the correspondence between connectivity-derived communities and conventional cellular anatomy.

Looking forward, xVDM points to a complementary path for spatial biology in which molecular sampling density and sequencing throughput become the determinants of resolution. This makes a different class of specimens approachable – large, opaque, or geometrically awkward tissues, embryos and tumors studied as whole intact units, and samples too fragile for serial sectioning – and it lets transcripts, proteins, lineage tags, and engineered perturbations share a single specimen and a single sequencing library. The harder problem sits at the inference layer. Across scales, the molecular network appears to carry information about tissue organization that is visible through molecular identity but not reducible to cell-state labels alone: sub-cellular rRNA partitioning emerged before any embedding, gene-level compactness tracked independently measured spatial autocorrelation, and recurring marker programs aligned with anatomical regions. Recovering these features from network structure and molecular phenotyping together is where we expect the next layer of biological interpretation to come from.

## Acknowledgments

We thank John Kilkus for his help with logistics, and Jing Li for helpful discussions. We thank Raghu Mirmira and Ryan Anderson for help procuring zebrafish embryos. We thank Marcus Clark for facilitating access to de-identified human tonsil specimens under University of Chicago IRB #22-1966. We thank Huili Wang for tissue embedding, cryosectioning, selection of germinal-center-containing tonsil sections, and preparation of the antibody-oligonucleotide probe cocktail used in the human tonsil experiments. We thank Bijentimala Keisham, Huili Wang, and Savas Tay for technical discussions and advice on adapting Prox-seq/SProx-seq antibody-oligonucleotide probes and protocols to the xVDM protein-target workflow, and the Tay laboratory for Prox-seq reagents and expertise. This work was supported by the Gordon and Betty Moore Foundation (GBMF9688), the National Institutes of Health (1R35GM143017-01), the National Science Foundation (2121044), and funding by the University of Chicago AI Initiative. Computational analyses were performed on multi-core Intel Xeon Gold 6248 CPU nodes maintained by the University of Chicago Research Computing Center.

## Author Contributions

N.Q. and R.Y. contributed equally to this work. J.A.W. conceived and supervised the study. J.A.W. and N.Q. designed, developed, and optimized the xVDM strategy. N.Q. performed zebrafish xVDM experiments and sequencing-library preparation, and contributed to data analysis and manuscript preparation. R.Y. developed and implemented the mammalian and protein-targeted xVDM workflows, performed mouse hippocampal chemistry optimization and human tonsil antibody-oligonucleotide xVDM experiments, generated the corresponding libraries, and contributed to analysis and interpretation of the protein-targeted data. M.Y. and H.C. contributed to computational analysis and visualization. J.A.W. developed the computational strategy and wrote the manuscript with input from all authors. All authors reviewed and approved the manuscript.

## Methods

### Circular DNA preparation

Single-stranded circular DNA Circ25-6G1 and Circ25-7G1 were prepared by splint-mediated ligation ^27^ (Fig. S1). Briefly, linear oligos 25.006G1 and 25.007G1 (20 µmol L^−1^) were annealed to their respective splints (splint6F5 and splint7F5, 100 µmol L^−1^) and subjected to iterative stepwise ligation with T4 DNA ligase (New England Biolabs, M0202L) under low Mg^2+^ and ATP conditions at 20 °C. Residual linear DNA was digested with 0.5 U µL^−1^ Exonuclease I (NEB, M0293L) and 5 U µL^−1^ Exonuclease III (NEB, M0206L) in 0.86× NEBuffer 1 at 37 °C for 45 min, followed by heat inactivation at 80 °C for 20 min. Circularized products were purified using Oligo Clean & Concentrator (Zymo Research, D4060), quantified by Nanodrop One Spectrophotometry (Thermo Scientific, 13-400-525), and assessed on a 15 % (w/v) TBE-Urea Gels (Invitrogen, EC68855BOX). Purified Circ25-6G1 and Circ25-7G1 were then diluted to 1 µmol L^−1^ and combined at a 1:1 molar ratio for subsequent circular DNA annealing.

### Zebrafish embryo collection

AB wild-type zebrafish (*Danio rerio*) were maintained and bred under standard conditions in accordance with protocols approved by the University of Chicago Institutional Animal Care and Use Committee (IACUC). To minimize maternal variation, embryos at 12, 18, and 24 hours post-fertilization (hpf) were collected from the same clutch. Embryos were enzymatically dechorionated with 1 mg mL^−1^ Pronase (Sigma-Aldrich, 10165921001) for 5–6 min at 28 °C, rinsed in embryo medium, and immediately fixed overnight at 4 °C in 4 % (w/v) Formaldehyde (Thermo Scientific, 28906) prepared in 1× PBS (Invitrogen, AM9624). Fixed embryos were dehydrated in 100 % methanol (Sigma-Aldrich, 34860-100ML-R) for 15–30 min at room temperature and stored at −80 °C for a minimum of 2 h before further processing.

### Experimental workflow of xVDM

A detailed schematic of the experimental workflow is shown in Fig. S2. Unless otherwise noted, all wash steps were performed for 5 min at room temperature.

#### Zebrafish embryo permeabilization

Before *in situ* reactions, methanol-stored embryos were rehydrated stepwise through 75 %, 50 %, and 25 % methanol in PBS (5 min each) and washed four times in 1× PBST (1× PBS supplemented with 0.1 % (v/v) Tween-20 (Sigma-Aldrich, P9416-100ML)). Permeabilization was performed with Thermolabile Proteinase K (New England Biolabs, P8111S) for 12 min at room temperature. Embryos 1, 2 (12 hpf) and 5, 6 (18 hpf) were treated at 2.5 × 10^−5^ U µL^−1^, whereas embryos 3, 4 (18 hpf) and 7, 8 (24 hpf) were treated at 5 × 10^−5^ U µL^−1^. The enzyme was inactivated by incubation at 55 °C for 15 min. Embryos were subsequently washed three times with 1× PBST.

#### *In situ* reverse transcription (RT)

To denature RNA secondary structures, permeabilized embryos were incubated in 1× PBS supplemented with 20 % (v/v) Formamide (Sigma-Aldrich, 47671-250ML-F), 0.5 U µL^−1^ Superase-In (Invitrogen, AM2696), 0.5 µg µL^−1^ rBSA (New England Biolabs, B9200S), and 4.4 mmol L^−1^ DTT at 4 °C for 1 h under gentle rotation (10 rpm), followed by 10 min at 65 °C and immediate cooling to 4 °C. After a water rinse, embryos were transferred into RT mix containing 1 µmol L^−1^ biotinylated poly(dT) primer (24.068Bio-30TVN), 400 µmol L^−1^ dNTP (Thermo Scientific, R0181), 0.5 µg µL^−1^ rBSA, 4.4 mmol L^−1^ DTT, 1 U µL^−1^ Superase-In, 10 U µL^−1^ Superscript III Reverse Transcriptase (Invitrogen, 18080085) in 1× First-Strand Buffer. The reaction was performed at 4 °C for 1 h under slow rotation (10 rpm), 60 °C for 3 min, and then 42 °C overnight on a nutating rocker (24 rpm). To eliminate excess primers and displaced cDNA after RT, embryos were treated with 1.43 U µL^−1^ Exonuclease I in 1× Exonuclease I Reaction Buffer at 4 °C for 1 h under slow rotation (10 rpm), followed by 1 h incubation at 37 °C. Embryos were washed three times with 1× PBST.

#### Cross-linking of rolling circle amplification (RCA) primers

A cross-linking mixture containing THIOCURE ETTMP 1300 (Bruno Bock, 345352-19-4), SM(PEG)12 (Thermo Scientific, A35398) and amine modified RCA primers (24.002 and 24.003) at a 2250:1 cross-linker-to-oligo ratio in 1× PBS was freshly prepared and incubated at room temperature for 30 min. After a rinse in 1× PBS, embryos were incubated in the mixture at room temperature for 1 h under gentle rotation (10 rpm). The reaction was terminated by quenching with 1 mol L^−1^ Tris pH 8 (Invitrogen, AM9855G) for 30 min at room temperature. After quenching, embryos were washed three times with 2× SSCT (2× SSC (Invitrogen, AM9770) supplemented with 0.1 % Tween-20).

#### Circular DNA annealing and RCA

Embryos were incubated overnight at 40 °C on a nutating rocker (24 rpm) in an annealing solution containing 100 nmol L^−1^ Circ25-6G1 and 100 nmol L^−1^ Circ25-7G1 in 1× hybridization buffer (2× SSC, 10 % formamide and 0.1 % Tween-20). The next day, embryos were washed in the 1× hybridization buffer at 40 °C for 30 min on a nutating rocker (24 rpm), and then sequentially washed with 2× SSCT, 1× SSCT (1× SSC supplemented with 0.1 % Tween-20) and 1× PBST.

Embryos were then rinsed once in water and incubated in an RCA mixture comprising 25 ng µL^−1^ T4 Gene 32 (New England Biolabs, M0300S), 250 µmol L^−1^ d(AUGC)TP, 0.5 µg µL^−1^ rBSA, 0.2 U µL^−1^ phi29 DNA Polymerase (New England Biolabs, M0269L) in 1× phi29 DNA Polymerase Reaction Buffer. For fluorescent labeling, Fluorescein-12-dUTP (Thermo Fisher Scientific, R0101) was added at a final concentration of 20 µmol L^−1^. The reaction was initiated with a 1 h incubation at 4 °C under gentle rotation (10 rpm) to allow reagent diffusion, followed by overnight incubation at 30 °C.

#### Bridging oligonucleotide annealing and UMI incorporation

After RCA, embryos were washed three times in 2× SSCT and transferred individually into fresh tubes. UEI oligo annealing was performed in UEI hybridization buffer (2× SSC, 5 % formamide and 0.1 % Tween-20) containing 50 nmol L^−1^ 21.004G1-BC, 50 nmol L^−1^ 21.004G2-BC, 100 nmol L^−1^ 21.073pt, and 100 nmol L^−1^ 21.074B by incubating at 4 °C for 1 h under slow rotation (10 rpm), followed by 2 h incubation at 50 °C. Embryos were then washed in UEI hybridization buffer for 1 h at 50 °C, followed by sequential washes with 2× SSCT, 1× SSCT and 1× PBST.

After a water rinse, embryos were transferred into a UMI incorporation mix including 1 mmol L^−1^ dNTP, 0.15 U µL^−1^ T4 DNA Polymerase (New England Biolabs, M0203L), 20 U µL^−1^ T4 DNA Ligase in 1× T4 DNA Ligase Reaction Buffer. Reactions were incubated at 4 °C for 1 h under slow rotation (10 rpm), followed by room temperature for 40 min. Embryos were then washed three times in 1× PBST.

#### Hydrogel-embedded *in vitro* transcription (IVT)

Embryos were transferred into a 96-well round-bottom untreated plate. IVT mixes were prepared by combining 2.1 µL water, 0.35 µL T7 promoter oligo (21.075, 10 µmol L^−1^), 0.35 µL T4 Gene 32 (10 µg µL^−1^), 1.75 µL USER Enzyme (1 U µL^−1^, New England Biolabs, M5505L), 10.5 µL rNTP (25 mmol L^−1^ each, New England Biolabs, N0466L), 3.5 µL T7 Enzyme Mix (Invitrogen, AM1334), 3.5 µL 10× Reaction Buffer and 1.05 µL THIOCURE ETTMP 1300 (213 µg µL^−1^). Following a rinse in water, 11.9 µL 4arm-PEG20K-Vinylsulfone (215 µg µL^−1^, Sigma-Aldrich, JKA7025-1G) was added to the IVT mixes, and 30 µL of the resulting reactions were immediately applied to individual embryos. The final concentrations of components in the 30 µL reactions were 100 nmol L^−1^ 21.075, 100 ng µL^−1^ T4 Gene 32, 0.05 U µL^−1^ USER Enzyme, 7.5 mmol L^−1^ rNTP mix, 10 % (v/v) T7 Enzyme Mix, 1× Reaction Buffer, 6.4 µg µL^−1^ THIOCURE ETTMP 1300, and 73.6 µg µL^−1^ 4arm-PEG20K-Vinylsulfone. Embryos were incubated for 2 h at room temperature, followed by 20 h incubation at 37 °C.

#### Nucleic acid purification

After IVT, hydrogels were denatured by adding 12 µL alkaline solution containing 42.5 mmol L^−1^ DTT (Thermo Scientific, P2325), 100 mmol L^−1^ EDTA pH 8.0 (Sigma-Aldrich, 03690-100ML), 457.5 mmol L^−1^ KOH (Honeywell, 319376-500ML), and incubating at 4 °C for 2 h. Reactions were neutralized with 12 µL acid solution composed of 0.6 mol L^−1^ Tris-HCl pH 7.5 (Fisher Scientific, BP1757-500) and 0.4 N HCl (Sigma-Aldrich, H9892-100ML). Samples were then supplemented with 30 µL of Proteinase K mix consisting of 0.28 % Tween-20, 0.09 U µL^−1^ Proteinase K (New England Biolabs, P8107S), 8.6 mmol L^−1^ Tris-HCl pH 7.5, to a final volume of 84 µL and incubated at 50 °C for 1 h. Nucleic acids (including IVT RNA and biotinylated genic cDNA) were purified using 1.8× RNAClean XP beads (Beckman Coulter, A63987) and eluted into 45 µL water.

#### Pulldown of biotinylated genic cDNA

20 µL Streptavidin beads (Invitrogen, 65001) were blocked with 50 µL blocking buffer containing 1 µg µL^−1^ rBSA, 1 µg µL^−1^ Salmon Sperm DNA (Invitrogen, AM9680) and 1 µg µL^−1^ Yeast tRNA (Invitrogen, AM7119) for 30 min at room temperature. Blocked beads were resuspended in 45 µL 2× BWT buffer (10 mmol L^−1^ Tris-HCl pH 7.5, 1 mmol L^−1^ EDTA, 2 mol L^−1^ NaCl, and 0.02 % Tween-20) and incubated with 45 µL RNAClean eluent from the previous step for 1 h at room temperature under slow rotation (20 rpm) to capture biotinylated genic cDNA. Bead-bound cDNA was stored at 4 °C until subsequent on-bead 3′ adapter ligation.

#### UEI cDNA synthesis

UEI RNA-containing supernatants from the biotin pull-down step were purified with 1.2× RNAClean XP beads and eluted in water. To remove residual DNA, samples were treated with 0.1 U µL^−1^ DNase I (New England Biolabs, M0303S) in 1× DNase I Reaction Buffer supplemented with 0.8 U µL^−1^ Superase-In for 30 min at 37 °C, followed by cleanup with 1.2× RNAClean XP beads and elution in water.

Purified UEI RNA was then reverse transcribed to cDNA in reactions containing 500 nmol L^−1^ RT primer (21.077), 500 µmol L^−1^ dNTP, 5 mmol L^−1^ DTT, 1 U µL^−1^ Superase-In, and 10 U µL^−1^ Superscript III Reverse Transcriptase in 1× First-Strand Buffer. RNA was first combined with 21.077 and dNTP, denatured at 65 °C for 5 min, and immediately cooled on ice. The remaining components were then added, and the reactions were incubated at 50 °C for 1 h. Reactions were terminated at 70 °C for 15 min and held at 4 °C. Residual RT primers were removed by adding 1.67 U µL^−1^ Exonuclease I and incubating at 37 °C for 30 min, followed by 20 min inactivation at 80 °C.

#### On-bead 3′ adapter ligation

To remove RNA, bead-bound genic cDNA was treated with 0.2 U µL^−1^ RNase H (New England Biolabs, M0297S) in 1× RNase H Reaction Buffer at 37 °C for 1 h. For adapter ligation, beads were resuspended in ligation mix containing 5 µmol L^−1^ 3′ adapter (24.004A), 1 mmol L^−1^ ATP, 25 % (w/v) PEG8000, and 1 U µL^−1^ T4 RNA Ligase 1 (New England Biolabs, M0204L) in 1× T4 RNA Ligase Reaction Buffer and incubated overnight at room temperature with gentle rotation (20 rpm). Biotinylated genic cDNA was then eluted by heating the beads in 95 % (v/v) formamide with 10 mmol L^−1^ EDTA at 95 °C for 10 min. The supernatants were purified with 1.5× Ampure XP beads (Beckman Coulter, A63881) and eluted in water.

#### PCR amplification and library pooling

Purified biotinylated genic cDNA was divided into four parallel PCR reactions per sample and amplified with Herculase II Fusion DNA Polymerase (Agilent, 600677). Each reaction contained 25 % of the cDNA, 300 nmol L^−1^ 21.046G1-BC, 300 nmol L^−1^ 21.081b, 400 µmol L^−1^ dNTP, and 2 % (v/v) Herculase II Fusion DNA Polymerase in 1× Herculase II Reaction Buffer. PCR cycling conditions were: 95 °C for 2 min; 5 cycles of 95 °C for 30 s, 56 °C for 30 s and 68 °C for 2 min; 13 cycles of 95 °C for 30 s and 68 °C for 2 min; and a final extension at 68 °C for 5 min before holding at 4 °C.

UEI cDNA (Exonuclease I-treated RT products) was amplified with Platinum Taq DNA HiFi Polymerase (Invitrogen, 11304029) in two parallel reactions per sample. Each reaction contained 5.2 % of the total UEI cDNA (corresponding to ∼1.27 % of the UEI RNA generated during IVT), together with 5 % (v/v) DMSO, 300 nmol L^−1^ 21.077-G1, 300 nmol L^−1^ 21.076BB, 3.3 µmol L^−1^ each of 4E4.interf1 and 4E.interf2, 200 µmol L^−1^ dNTP, 2 mmol L^−1^ MgSO_4_, and 0.02 U µL^−1^ Platinum Taq DNA HiFi Polymerase in 1× High Fidelity Buffer. 4E4.interf1 and 4E.interf2 served as interference oligos to prevent PCR recombination of intermediate UEIs. PCR cycling conditions were: 95 °C for 2 min; 1 cycle of 95 °C for 30 s, 66 °C for 30 s, and 68 °C for 2 min; 18 cycles of 95 °C for 30 s and 68 °C for 2 min; and a final extension at 68 °C for 5 min before holding at 4 °C.

All PCR products were purified with 0.75× Ampure XP beads and eluted in 10 mmol L^−1^ Tris-HCl pH 8.0 (Fisher Scientific, BP1758-500). Libraries were pooled and sequenced on a NextSeq 500 (Illumina) or Aviti (Element) instrument.

### Thiol-anchored xVDM for zebrafish and human tonsil

To test whether matrix seeding for xVDM could be achieved by an orthogonal chemistry, fixed 24 hpf zebrafish embryos were processed as above through rehydration, permeabilization, and pre-RT incubation, and then assigned to comparison conditions that differed only in the chemistry used to anchor RCA primers to the fixed specimen matrix. These conditions included a no-circular-template control, a standard amine-coupling condition, a low-pH-to-normal-pH amine-coupling condition, and TCEP-dependent thiol-oriented conditions. For thiol-oriented conditions, thiol-bearing oligos were chemically reduced immediately before use. Following cross-linking and quenching, circular DNA templates were annealed overnight at 40 °C, and rolling-circle amplification was performed with phi29 DNA polymerase in the presence of fluorescein-12-dUTP. Embryos were then washed for fluorescence imaging.

Whole-mount embryos were imaged at 10× and 40× magnification using fixed acquisition settings within each comparison set. Displayed panels were used as qualitative comparisons of RCA-derived fluorescence across anchoring conditions.

### Fluorescence-based chemistry optimization in mouse hippocampal sections

To bridge the whole-mount zebrafish fluorescence experiments and the final human tonsil sequencing workflow, we performed fluorescence-only optimization of matrix-seeding chemistry in thick floating mouse hippocampal sections. These mammalian-tissue experiments were used to compare cross-linking chemistries by fluorescent rolling-circle-amplification (RCA) readout and were not carried through UEI generation, hydrogel IVT, or sequencing.

#### Mouse hippocampal floating-section preparation and permeabilization

All animal experiments and procedures were performed in accordance with the University of Chicago Institutional Animal Care and Use Committee (IACUC) protocols. C57BL/6 mouse (male, 8 weeks of age) was transcardially perfused with ice cold PBS followed by 4 % PFA. The collected hippocampus hemispheres were then post fixed in 4 % PFA at 4 °C for 20 h. The hippocampal hemispheres were then cryopreserved in 15 % sucrose followed by 30 % sucrose until they sank. The tissue was then embedded in O.C.T. and stored at −80 °C. The tissue was then sectioned into 400 µm sections using a cryostat (Thermo NX50). Mouse hippocampal sections (400 µm) were collected directly into cold 1× PBS supplemented with 1 mmol L^−1^ EDTA and dehydrated through 25 %, 50 %, 75 %, and 100 % methanol (5 min each), followed by storage at −80 °C. Before use, sections were rehydrated through 75 %, 50 %, and 25 % methanol in PBS (5 min each) and washed four times in 1× PBST. Sections were permeabilized with thermolabile Proteinase K diluted 1:2400 in PBST for 15 min at 23 °C to 24 °C, rinsed once in PBST, incubated in PBST at 55 °C for 15 min, washed three times in PBST, and maintained at 4 °C until chemistry optimization.

#### Pre-annealing conditioning and cross-linking comparison

Sections were incubated in a pre-RT conditioning buffer containing 20 % formamide, 1.4× PBS, 4.4 mmol L^−1^ DTT, 0.5 mg mL^−1^ BSA, and 0.5 U µL^−1^ Superase-In for 1 h at 4 °C, followed by 15 min at 65 °C and return to 4 °C. Sections were then assigned to fluorescence comparison conditions including a no-circular-template negative control, a standard amine-coupling condition at near-neutral pH, a pH-switched amine condition in which samples were first exposed to a low-pH diffusion step and then to a higher-pH reaction step, and a thiol-directed condition. Where indicated in the fluorescence comparison panels, additional amine-control variants included overnight 0.5 % Triton X-100 pretreatment or 10 % PEG8000 supplementation.

Amine-based conditions used freshly prepared mixtures of THIOCURE ETTMP 1300, SM(PEG)12, and the 5′-amine-modified RCA primers 24.002 and 24.003 at a 2250:1 cross-linker-to-oligo ratio. Cross-linking mixes were pre-incubated for 30 min at room temperature before tissue application. For pH-staged amine conditions, sections were first incubated in reaction mix adjusted with MES (final 0.1 mol L^−1^, pH 6.1) at 4 °C to favor diffusion, then transferred to a second aliquot adjusted with borate (final 0.05 mol L^−1^, pH 7.9) or PBS and incubated at room temperature to drive coupling. Thiol-directed conditions used TCEP-reduced thiol-modified versions of 24.002 and 24.003 together with 4arm-PEG2K-NH_2_ and SM(PEG)12 at the same 2250:1 ratio. Thiol-modified oligos were reduced immediately before use with 10 mmol L^−1^ TCEP for 2 h at room temperature, and the tissue reaction was carried out using a low-pH diffusion step followed by a room-temperature reaction step in the presence of 10 mmol L^−1^ TCEP. Cross-linking reactions were quenched, and sections were washed three times in 2× SSCT before circular-template annealing.

#### Circular DNA annealing, fluorescent RCA, and imaging of mouse sections

Paired circular DNA templates (6G1 and 7G1; 100 nmol L^−1^ each) were annealed overnight at 40 °C in hybridization buffer containing 2× SSC, 10 % formamide, and 0.1 % Tween-20. Sections were then washed for 30 min at 40 °C in diluted hybridization buffer, followed by sequential washes in 2× SSCT, 1× SSCT, and PBST. After a water rinse, RCA was performed in 1× phi29 DNA polymerase reaction buffer containing 20 µmol L^−1^ Fluorescein-12-dUTP, 25 ng µL^−1^ T4 gene 32 protein, 250 µmol L^−1^ d(AUGC)TP, 0.5 mg mL^−1^ BSA, and 0.2 U µL^−1^ phi29 DNA polymerase. Sections were incubated for 1 h at 4 °C with rotation to allow reagent diffusion and then overnight at 30 °C without rotation. After washing in 2× SSCT and PBST, sections were transferred to a flat-bottom imaging dish and imaged; acquisition settings were held constant within each comparison set. Fluorescence was used as a qualitative readout of circular-template-dependent nanoball formation in mammalian tissue.

These mouse-section fluorescence experiments informed selection of the mammalian-tissue thiol/TCEP cross-linking strategy later used for human tonsil xVDM, but the mouse sections themselves were not advanced to library preparation or sequencing.

#### Human tonsil tissue preparation and antibody-oligonucleotide staining

De-identified human tonsils were obtained from University of Chicago Medicine after routine tonsillectomies according to institutional guidelines. This study was approved by The University of Chicago Institutional Review Board (IRB# 22-1966). The tonsil samples were embedded in O.C.T. compound, stored at −80 °C, cryosectioned, and mounted on Fisherbrand Superfrost Plus Microscope Slides. Tonsil cryosections were fixed in freshly prepared 1 % PFA for 10 minutes and placed in a ProPlate Multiwell chamber (Grace Bio-Labs). The antibody inputs used to prepare antibody-oligonucleotide conjugates (AOCs) were purified anti-human IgD antibody (BioLegend, 348202, RRID:AB_10550095), purified anti-human CD9 antibody (BioLegend, 312102, RRID:AB_314907), purified anti-human CD3 antibody (BioLegend, 300302, RRID:AB_314038), and purified anti-human CD24 antibody (BioLegend, 311102, RRID:AB_314851). Sections were blocked and incubated with AOCs (2.5 nmol L^−1^ each) in binding buffer (0.2 % BSA, 0.2 mg mL^−1^ sonicated salmon sperm DNA, 100 µg mL^−1^ mouse isotype antibodies) at 4 °C for 1.5 h. After washing, bound AOC oligos were subjected to splint ligation using 9.5 nmol L^−1^ splint oligo (25.012), 10 U µL^−1^ T4 DNA ligase, and 1× T4 DNA Ligase Reaction Buffer at 37 °C for 30 min.

Sections were then treated with 0.5 U µL^−1^ Quick CIP phosphatase, 1.43 U µL^−1^ Exonuclease I in 1× rCutSmart Buffer at 37 °C for 30 min to digest unligated AOC oligos. Tonsil specimen 2 received both enzyme treatments and Tonsil specimen 1 received neither. Cross-linking of matrix anchored oligos used TCEP-reduced thiol-modified 24.002 and 24.003 together with 4arm-PEG2K-NH_2_ and SM(PEG)12 at a 2250:1 ratio. The cross-linking mix was pre-incubated for 30 min at room temperature before the tissue reaction in 5 mmol L^−1^ TCEP. Following cross-linking of matrix-anchored oligos, paired circular DNA templates (6G1 and 7G1; 100 nmol L^−1^ each) and biotinylated primer (25.015 at 1 µmol L^−1^) were annealed overnight at 40 °C in hybridization buffer containing 2× SSC, 10 % formamide, and 0.1 % Tween-20. Sections were then washed for 30 min at 40 °C in diluted hybridization buffer, followed by sequential washes in 2× SSCT, 1× SSCT, and PBST. Sections were then incubated with 0.2 U µL^−1^ DNA Polymerase I, 1 mmol L^−1^ dNTP in 1× NEBuffer 2 at 23 °C to 24 °C for 45 min followed by an Exonuclease I treatment to remove residual primers. RCA was then performed as described above.

#### PCR amplification and library pooling of protein-target libraries

The purified biotinylated proximity ligation product was amplified with Platinum Taq DNA HiFi Polymerase (Invitrogen, 11304029) in two parallel reactions per sample. Each reaction contained 50 % of the total proximity ligation product, 300 nmol L^−1^ 21.046G1-BC, 300 nmol L^−1^ 25.018, 200 µmol L^−1^ dNTP, 2 mmol L^−1^ MgSO_4_, and 0.02 U µL^−1^ Platinum Taq DNA HiFi Polymerase in 1× High Fidelity Buffer. PCR cycling conditions were: 95 °C for 2 min ; 5 cycles of 95 °C for 30 s, 50 °C for 30 s, and 68 °C for 2 min; 20 cycles of 95 °C for 30 s and 68 °C for 2 min; and a final extension at 68 °C for 5 min before holding at 4 °C.

The PCR products were purified with 0.9× Ampure XP beads and eluted in 10 mmol L^−1^ Tris-HCl pH 8.0 (Fisher Scientific, BP1758-500). Libraries were pooled and sequenced on an Aviti (Element) instrument.

### Sequencing data processing

Sequencing reads were parsed according to the read layouts and sample barcodes in Table S1. UMI type I, UMI type II, UEI, and cDNA insert sequences were clustered with EASL ^9;14^ using a one base edit rule. UMI and UEI clusters were removed when one base accounted for at least 75% of the positions in the sequence.

For UEI libraries, each UEI was assigned to the UMI pair with the greatest read support. Unless a different value was specified in the sample settings file, retained UMI pairs required at least two reads and retained UMIs required at least two linked UEIs. The largest connected component of the retained UMI graph was used for GSE embedding.

For cDNA libraries, reads assigned to the same UMI cluster were split into insert sub consensus groups by the first ten bases of the insert. Within each group, the consensus base at each position was the strict majority base. Ties were written as N. Insert sequences were trimmed at sample specific termination sequences, and inserts below the minimum accepted length were removed.

Retained cDNA sub consensuses were aligned with STAR to the *Danio rerio* GRCz11 reference index built from Ensembl release 109 annotations. Alignments were filtered to remove reads with no matched bases or long homopolymers. For each retained alignment, overlaps with the GTF file were used to assign contig, gene name, gene id, biotype, and transcript. rRNA and mitochondrial rRNA calls were given priority when present.

After UMI clustering, cDNA UMI clusters were matched to UEI UMI clusters by their clustered UMI sequence. For each matched UEI node, the analysis file stores all assigned cDNA sub consensus calls as aligned lists of starts, mutations, contigs, genes, biotypes, transcripts, sub consensus read counts, query names, and UEI read counts. These files provided the input for hub analysis, cell annotation, and gene level compactness analysis.

### Feature matrices

rRNA and mitochondrial rRNA calls were collapsed to rRNA and Mt_rRNA before this support was tallied. The hub and cell expression matrices used sub consensus read counts as feature counts. Unless stated otherwise, sub consensuses with more than one distinct gene call were excluded from these matrices. Repeated calls to the same gene within one hub were counted once. Unannotated genome calls were stored as contig features and were not mixed with gene features. This rule separates the rarefaction gene support metric from the expression matrices used for hub, cell, and compactness analyses.

## Data availability

Raw sequencing data generated in this study have been deposited in the NCBI SRA under accession PRJNA1472691. The transcript anchored VDM control data from ^14^ are available under PRJNA1004618. The Stereo-seq reference data were obtained from ^23^.

## Code availability

The code used for read parsing, UMI and UEI clustering, cDNA consensus calling, graph construction, GSE embedding, segmentation, hub analysis, protein target analysis, and reference registration is available at https://github.com/wlab-bio/xVDM. The repository includes the sample settings files, figure scripts, and software environment file. No restrictions apply beyond the license stated in the repository.

## Materials availability

All oligonucleotide sequences are listed in Table S1. Commercial antibody inputs used for the human tonsil antibody-oligonucleotide panel are listed in *Methods: Human tonsil tissue preparation and antibody-oligonucleotide staining*.

## Supplementary Information

**Figure S1.**
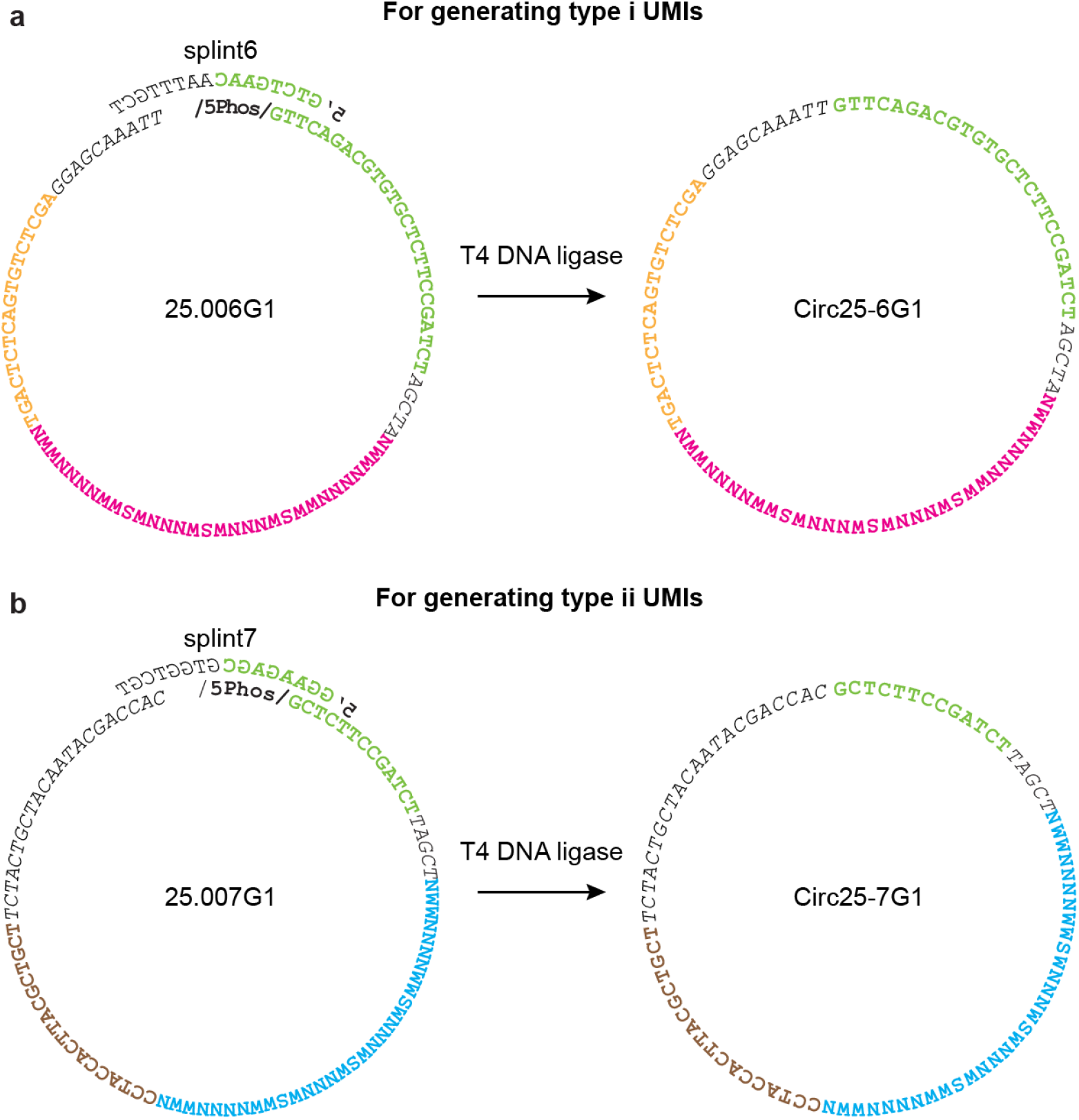
Circular DNA preparation. **a**, For generating “type i” UMIs, 8 bases at the 5′ and 3′ ends of splint6 anneal to 25.006G1, bringing its termini together for T4 DNA ligation. **b**, For generating “type ii” UMIs, 8 bases at the 5′ and 3′ ends of splint7 anneal to 25.007G1, bringing its termini together for T4 DNA ligation.

**Figure S2.**
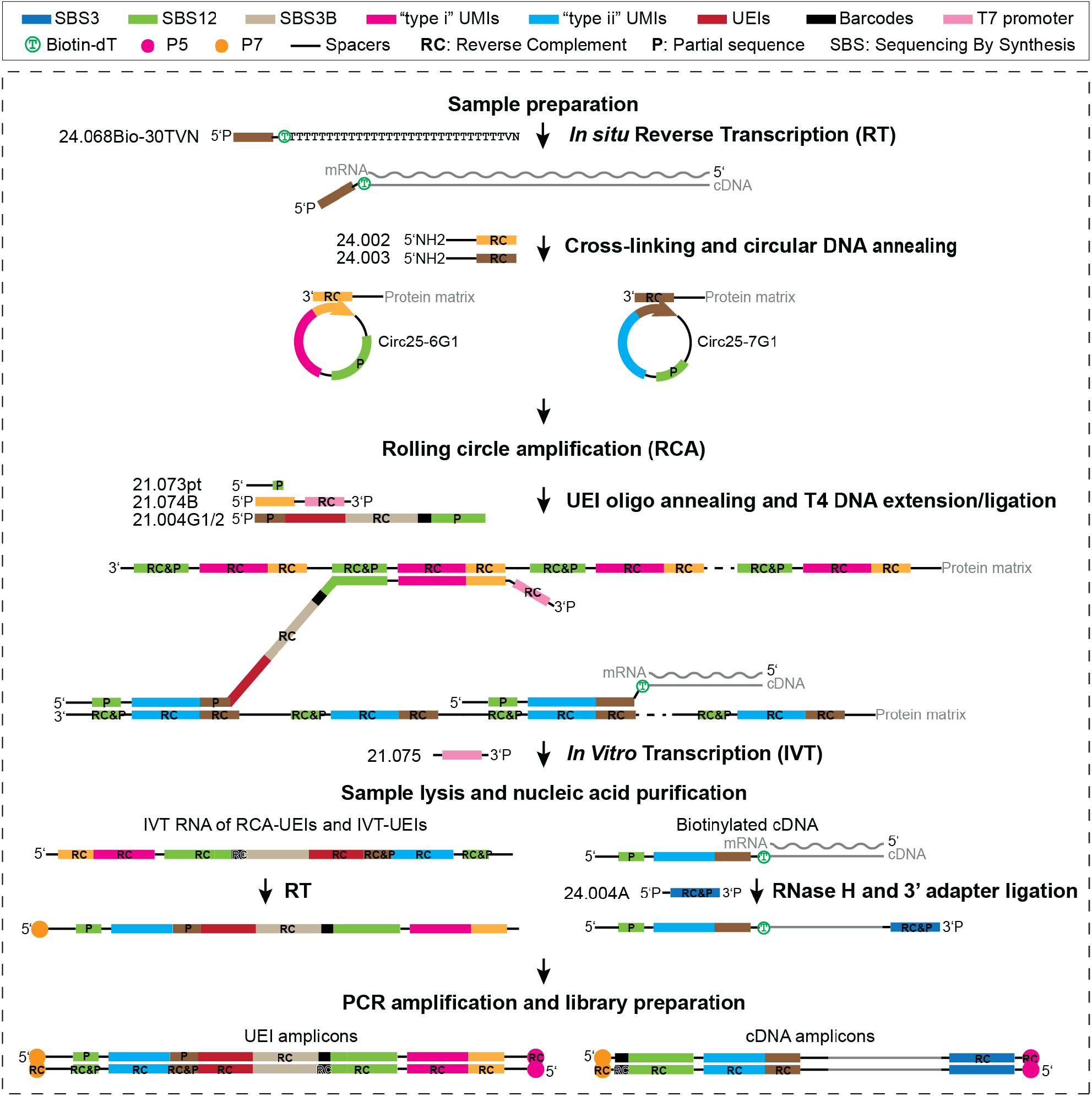
Overview of the experimental workflow of xVDM. Green sequences are not identical; they represent SBS12, distinct partial fragments of SBS12 (P), or its reverse complement (RC). The green portion of 21.073pt anneals specifically to DNA nanoballs generated by Circ25-7G1, while the green portion of 21.004G1/2-BC anneals only to those generated by Circ25-6G1. This staggered design provides efficient access to UMI sequences in both UEI and cDNA amplicons without reading through long spacer regions during sequencing. For complete sequence information, refer to Table S1

**Figure S3.**
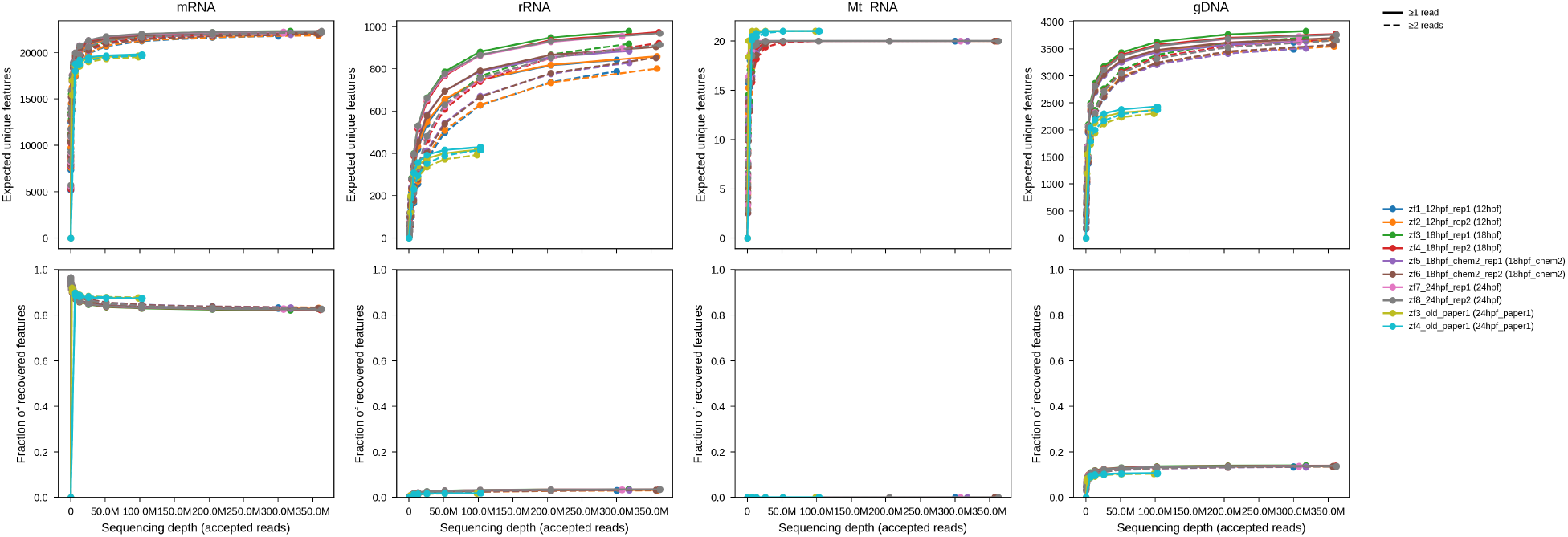
Rarefaction by feature type. Number of mRNA/rRNA/MT-rRNA/non-genic sequence inserts in cDNA libraries as a function of read depth (top row). Fraction of total features (bottom row). Solid lines denote features supported by 1-read sub-consensuses whereas dotted lines show features supported by at least 2 reads.

**Figure S4.**
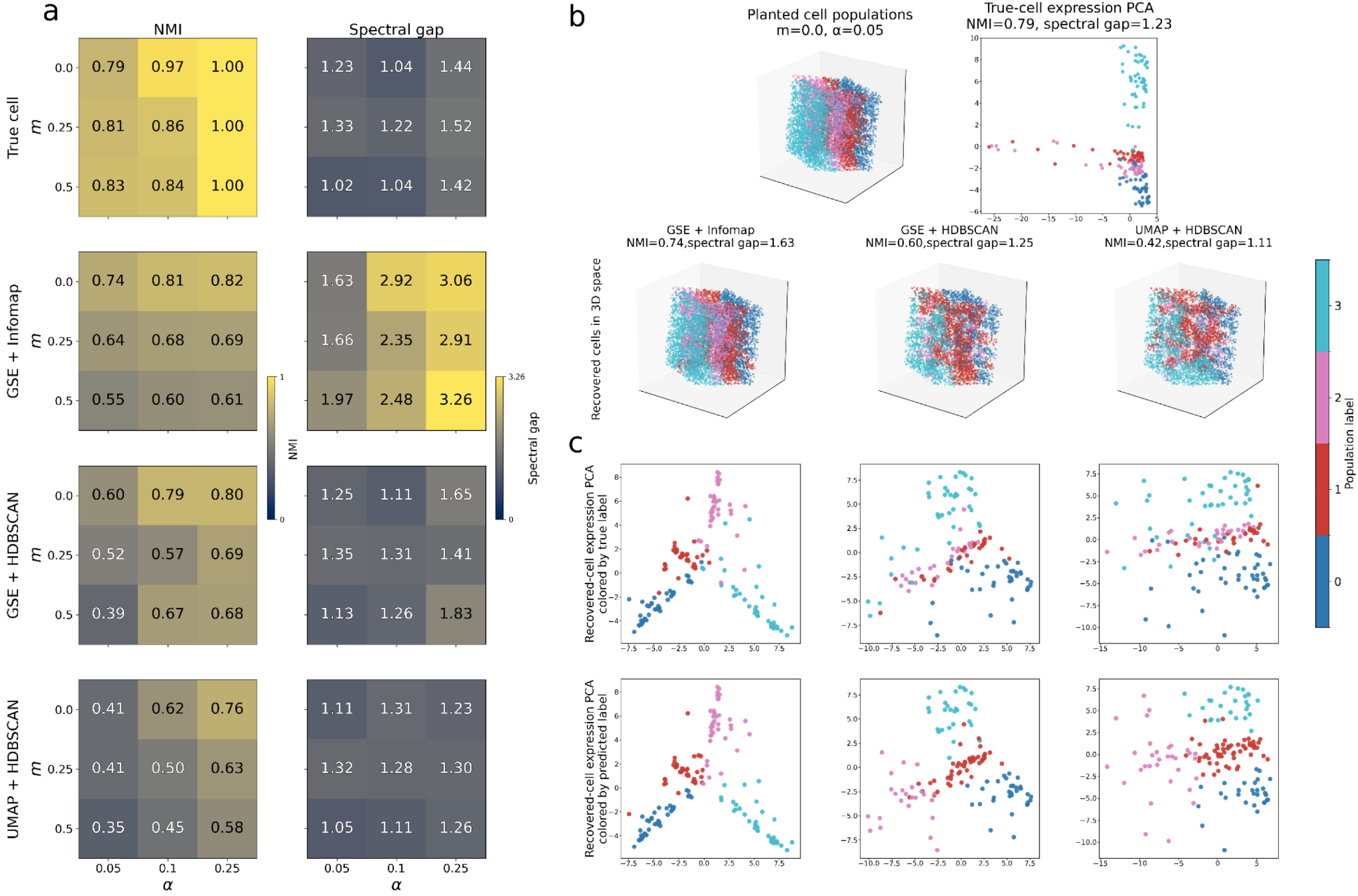
Recovering planted cell-populations in simulation. Cells were assigned evenly to four planted populations. A spatial mixing parameter, 0 ≤ *m* ≤ 1, controlled the spatial organization of population labels, with *m* = 0 corresponding to spatially coherent and lineage-like populations and *m* = 1 corresponding to population labels being spatially well-mixed. A marker-strength parameter, 0 ≤ *α* ≤ 1, controlled the marker genes to all genes transcript abundance ratio. Each population was assigned a small, orthogonal marker-gene program. **a**, Normalized mutual information (NMI) and expression spectral gap (PC3/PC4) for recovery of four planted cell populations sweeping parameter *m* and *α*. NMI was computed between the planted population label of each UMI and the predicted population label assigned to that UMI through true-cell or recovered-cell aggregation, with higher values indicating stronger agreement. Results compare the recovery upper bound (true Voronoi cell aggregation) with recovered cells obtained by GSE+Infomap, GSE+HDBSCAN, and UMAP+HDBSCAN. **b**, Representative simulation condition (*m* = 0, *α* = 0.05), showing planted populations, true-cell expression PCA, and recovered cells in 3D space. **c**, Recovered-cell expression PCA for the same condition, colored by true population labels (the population label that majority UMIs inherit from the true cell assignment) or by predicted labels from k-means clustering in expression space.

**Figure S5.**
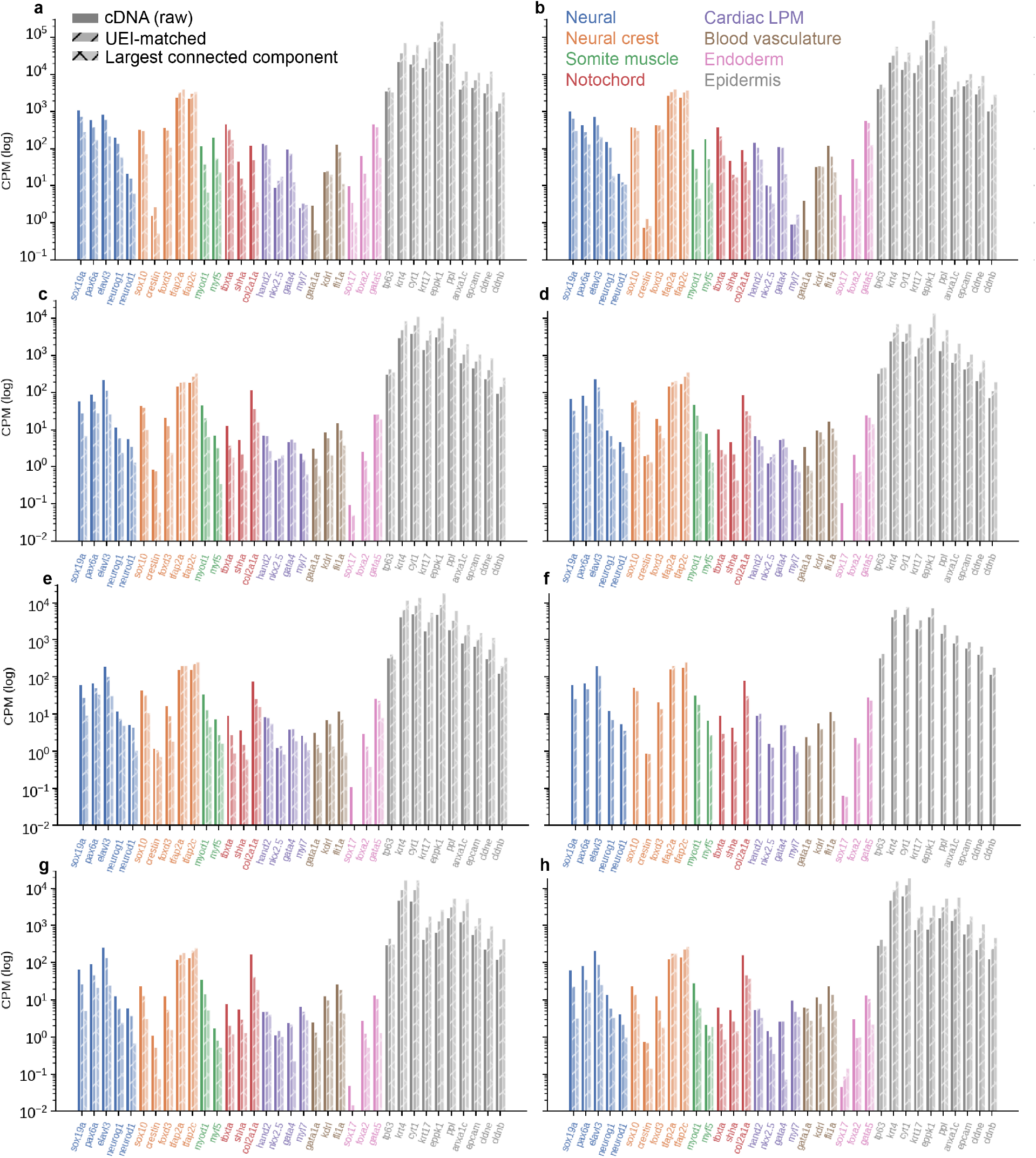
Retention of zebrafish development marker genes across different levels of data analysis. For each zebrafish specimen, canonical marker-gene CPM is tracked across the cDNA-library to UEI-library matching process, and then to their representation within the largest connected UEI-network, used for GSE-embedding. Values are normalized within each stage as counts per million. **a**-**h** corresponds to zf1 through zf8, respectively.

**Figure S6.**
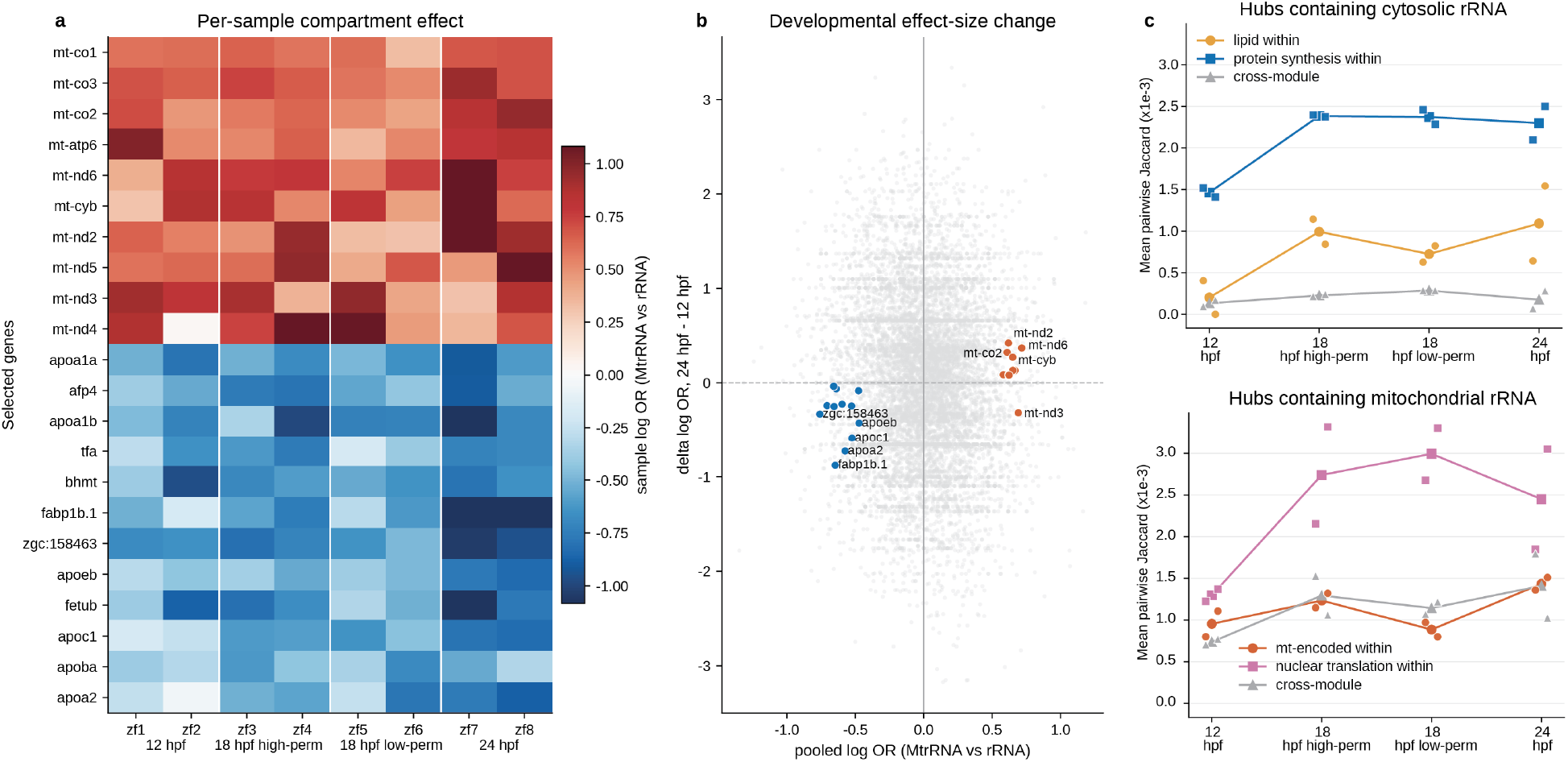
cDNA hub compartment analysis. **(a)** cDNA-hub compartment analysis. Within each embryo, hubs carrying protein-coding signal with cytosolic rRNA but no mitochondrial rRNA were compared with hubs carrying protein-coding signal with mitochondrial rRNA but no cytosolic rRNA after balancing for UMI type and hub complexity. Hub complexity was the total number of detected features on the hub, capped at a six-or-more bin. Points are protein-coding genes. The x-axis gives the inverse-variance fixed-effect log odds ratio for mitochondrial-rRNA-containing hubs relative to cytosolic-rRNA-containing hubs. (**b**) Developmental change in the same compartment effect. Each point is a gene. The x-axis shows the pooled fixed-effect log odds ratio from Fig. 3a. The y-axis shows the 24 hpf fixed-effect stage estimate minus the 12 hpf fixed-effect stage estimate. This panel summarizes effect-size change and was not used as a separate hypothesis test. (**c**) Module signals in the same matched hub sets across developmental groups. Points show embryos and lines show group means. The y-axis is mean pairwise Jaccard multiplied by 10^3^. The left plot shows hubs with cytosolic rRNA. The right plot shows hubs with mitochondrial rRNA. Lipid, protein synthesis, mitochondrial, and nuclear translation modules were set before this comparison.

**Figure S7.**
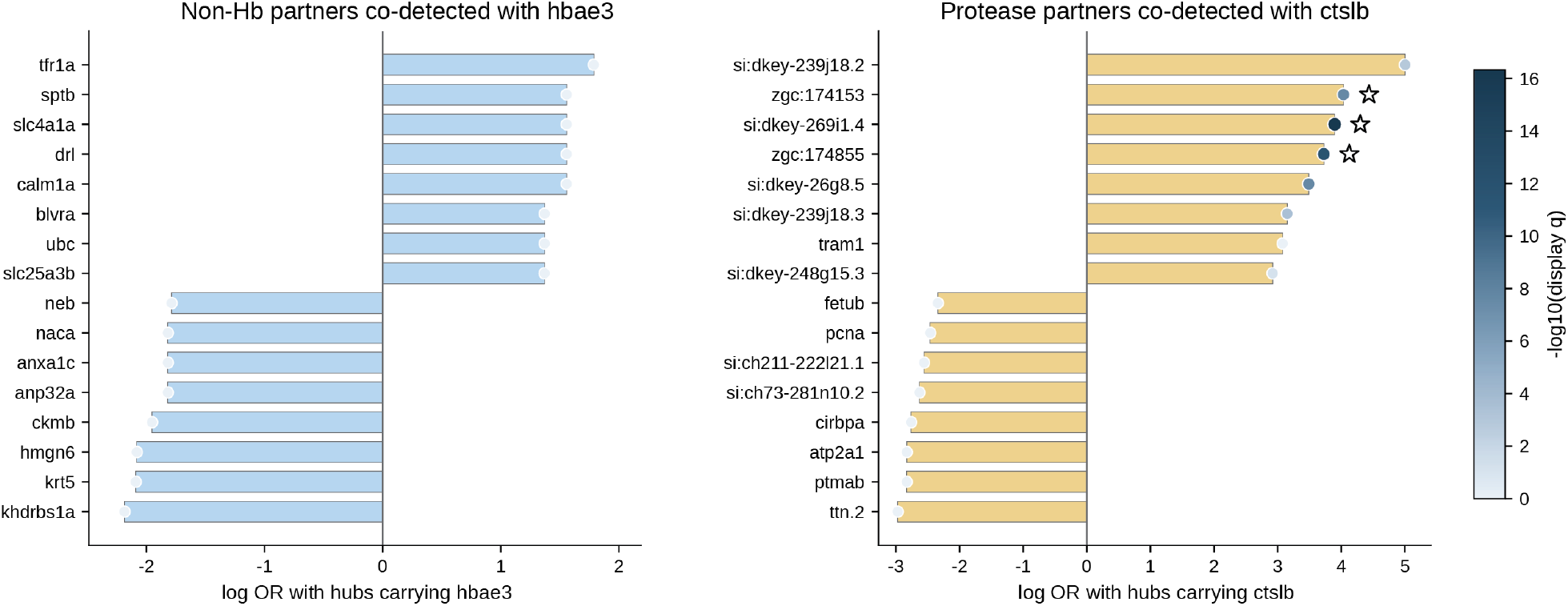
Matched 24 hpf cDNA hub analysis for *hbae3* and *ctslb*. For each analysis, the named gene was used as an anchor only in the statistical sense. Hubs carrying that gene were matched within each 24 hpf embryo to hubs lacking that gene by UMI type and detected feature count. UMI type denotes the type i or type ii RCA UMI template used to generate the hub. Each bar shows the fixed-effect log odds ratio for the listed gene in anchor-carrying hubs relative to matched hubs lacking the anchor. Dot size and color show fixed-effect *q* values after Benjamini–Hochberg adjustment over tested protein-coding genes. **a**, The *hbae3* anchor produced no clear non-hemoglobin gene set. **b**, The *ctslb* anchor recovered genes from the protease module, including *si:dkey-269i1*.*4, zgc:174855, zgc:174153*, and *zgc:158463*. Stars mark genes specified before the test.

**Figure S8.**
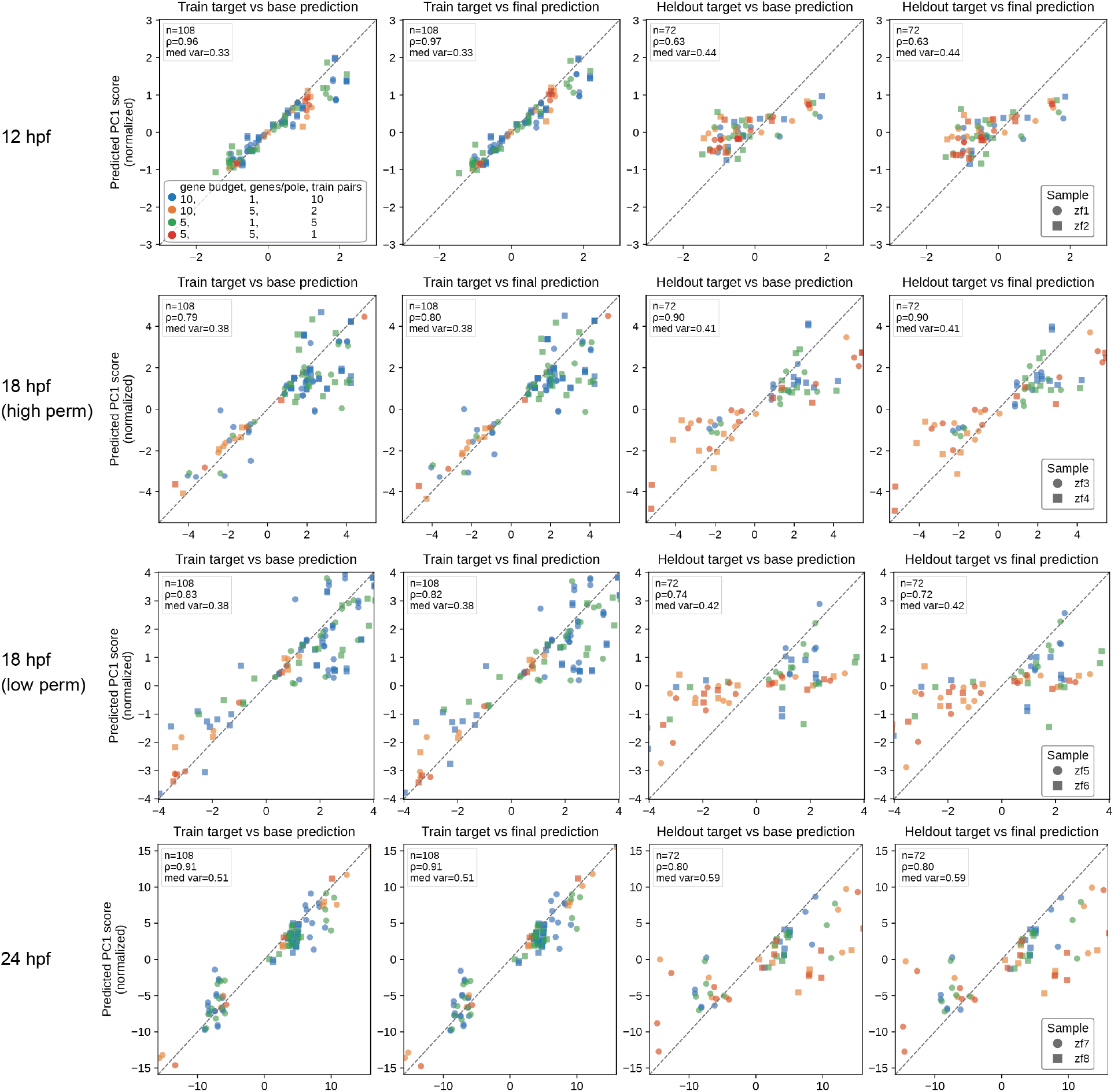
Train and held-out marker-field correlograms for reference-guided registration. PC1 summaries of target-prediction agreement are shown for the held-out registration benchmark. Within each run, rank-rescaled training and held-out pole-pair fields were projected into a target-defined PC basis. Only PC1 is shown. Columns compare training/base, training/final, held-out/base, and held-out/final predictions, where “base” denotes the feature-only sparse-transport map and “final” denotes the connectivity-refined map. The benchmark swept training marker budgets of five and ten genes per side and genes-per-pole regimes of one and five, using three folds and three held-out pole pairs per run; held-out scoring used single-gene pole pairs disjoint from the fitted markers. Rows show embryo pairs by stage: specimens 1 and 2, 12 hpf; specimens 3 and 4, 18 hpf high permeabilization; specimens 5 and 6, 18 hpf low permeabilization; and specimens 7 and 8, 24 hpf. Agreement in the held-out/final panels provides a compact check for transfer beyond the marker axes used to fit each map.

**Figure S9.**
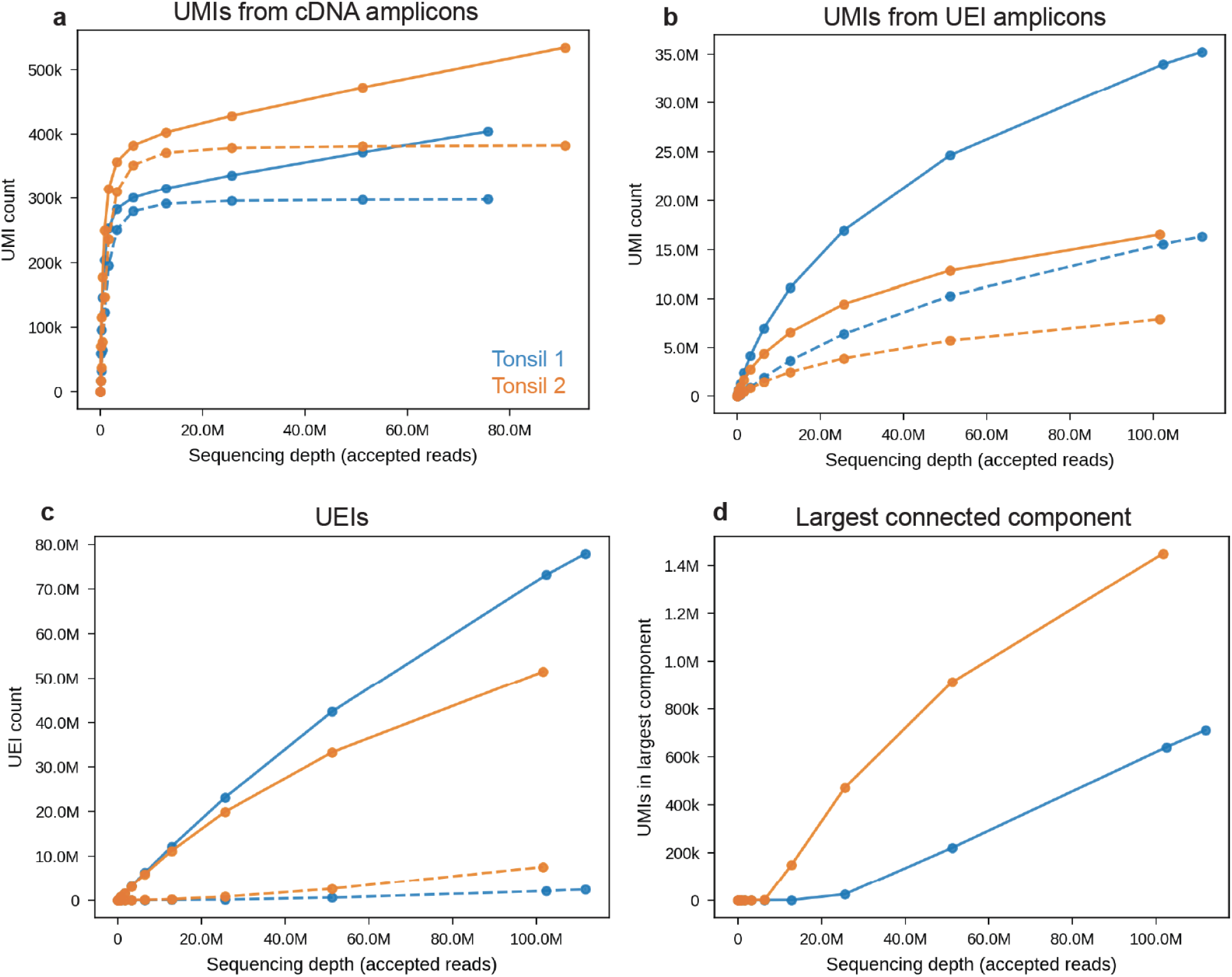
Protein-target xVDM proof of principle in human tonsil. **a–d**, Rarefaction of antibody-oligonucleotide-derived xVDM libraries from human tonsil, shown in the same style as transcript rarefaction in Fig. 2l–o. These data establish compatibility of matrix-seeded xVDM chemistry with antibody-oligonucleotide conjugates

**Figure S10.**
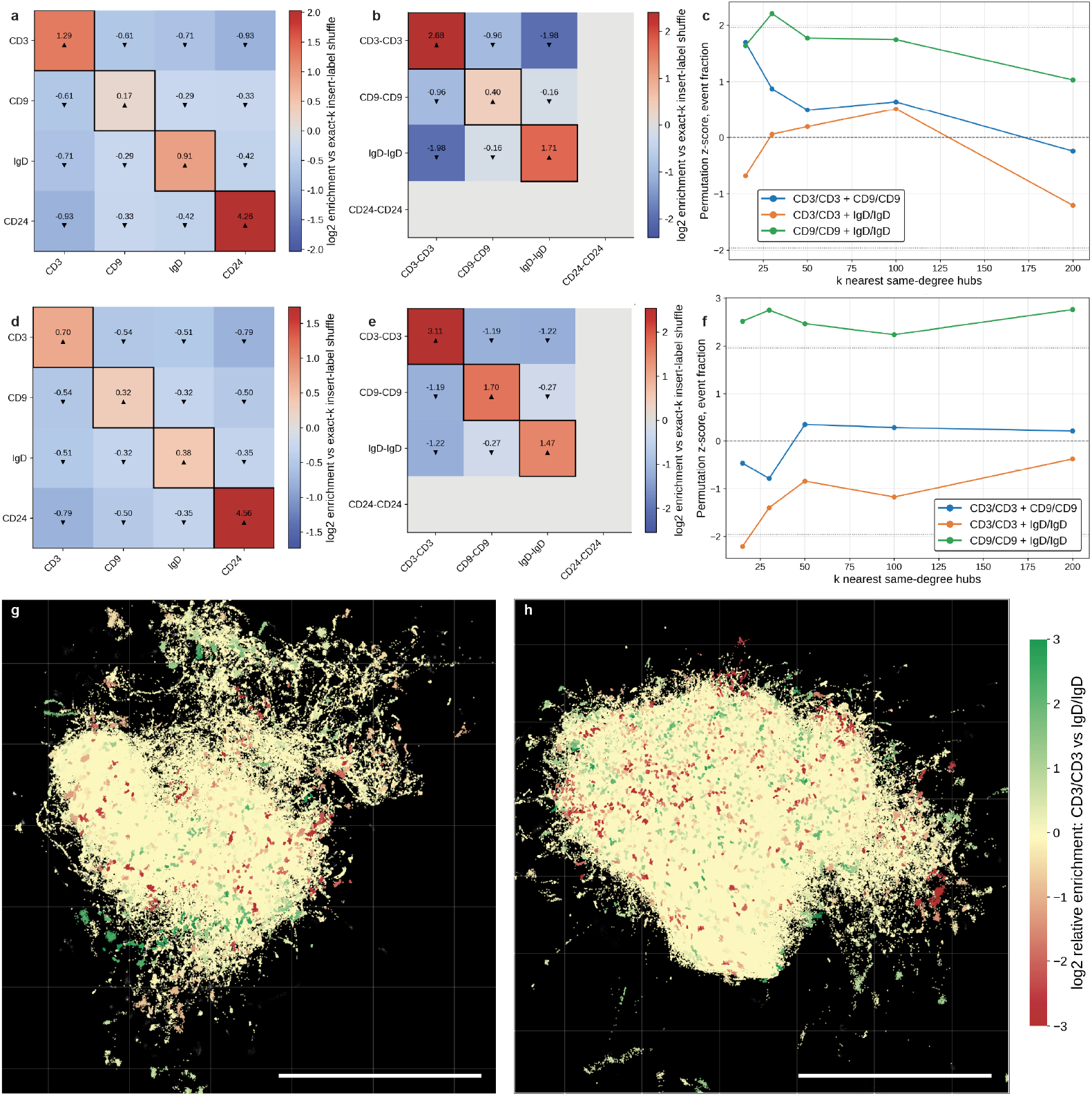
Protein xVDM captures local protein context and stratification of B and T cell regions in human tonsil. **(a**,**d)** Different-insert epitope co-localization on cDNA-library hubs for tonsil specimen 1 (**a**) and tonsil specimen 2 (**d**). A qualifying insert is an antibody-derived insert containing exactly two recognized epitope motifs. These panels use hubs with exactly two qualifying inserts. Color shows log_2_ enrichment, where log_2_ denotes the base-2 logarithm, relative to permutations that shuffle epitope-pair labels among the two fixed insert slots per hub while preserving the number of hubs and the global abundance of each epitope-pair label. Triangles mark heat-map cells passing Benjamini–Hochberg false-discovery-rate control at *q* ≤ 0.05, where *q* is the adjusted *P* value for enrichment or depletion. A heat-map cell is counted only when the compared epitopes can be assigned to two distinct inserts on the same hub. Same-epitope comparisons, including CD3 with CD3 and IgD with IgD, are enriched in both tonsils. **(b**,**e)** The same exact two-insert hub stratum restricted to homo-epitope insert labels for tonsil specimen 1 (**b**) and tonsil specimen 2 (**e**). A homo-epitope insert label, such as CD3/CD3 or IgD/IgD, denotes an insert whose two recognized motifs come from the same epitope class. **(c**,**f)** UEI-based *K*-nearest-neighbor co-localization for tonsil specimen 1 (**c**) and tonsil specimen 2 (**f**), restricted to hubs with exactly one qualifying protein insert. Here *K* is the number of nearest hubs within the same exact qualifying-insert-count stratum. The plotted *z*-score is the observed pair-label event fraction minus the permutation mean, divided by the permutation standard deviation. CD3/CD3 and IgD/IgD signals separate across neighborhoods, consistent with T cell and B cell zones. **(g**,**h)** GSE maps for tonsil specimen 1 (**g**) and tonsil specimen 2 (**h**). Hubs are colored by Infomap-cluster-level log_2_ relative enrichment of CD3/CD3 versus IgD/IgD, computed from *K* = 30 nearest-neighbor neighborhoods within the exact one-insert hub stratum. CD3-rich and IgD-rich regions occupy distinct parts of the inferred tonsil maps. Scale bars span 25 diffusion units.

**Table S1.**
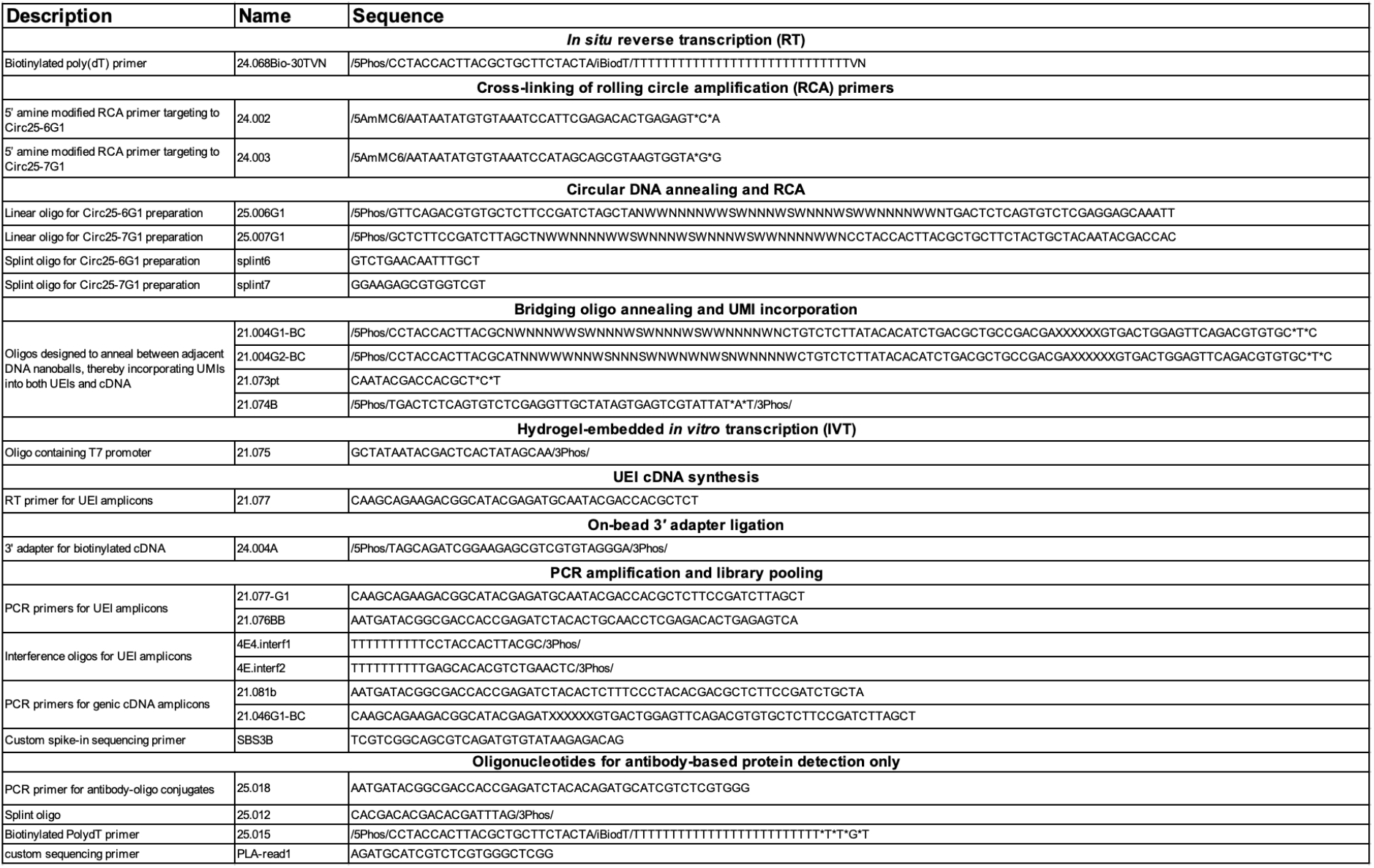
Oligonucleotides used in this study. /iBiodT/ denotes an internal Biotin dT. /5AmMC6/ designates a 5′ amine modification. Terminal phosphate groups are indicated as /5Phos/ (5′ end) and /3Phos/ (3′ end). Phosphorothioate linkages are represented by an asterisk (*). The sequence placeholder “XXXXXX” denotes sample-specific 6-nt barcodes. SBS3B is a custom sequencing primer for read 1 of UEI amplicons; in some sequencing kits it is already included, and no additional spike-in is required.

**Table S2.** Observed full-depth library summary across developmental conditions. Values are medians across embryos, with ranges in parentheses. The 24 hpf control row is shown only as full-depth context; the like-to-like chemistry comparison is given in Table S3. The cDNA-hub values are library-yield counts, not the row counts of the filtered matrices used for hub co-detection.

| *(a) UEI libraries* |  |  |  |  |
| --- | --- | --- | --- | --- |
| Condition | *n* | Accepted reads<sup>1</sup>(M) | Mol. nodes<sup>2</sup> (M) | Mol. nodes $\geq 2$ reads<sup>3</sup>(M) |
| xVDM, 12 hpf | 2 | 408.1 (388.0–428.1) | 66.4 (63.7–69.2) | 39.8 (38.2–41.4) |
| xVDM, 18 hpf high perm. | 2 | 416.5 (399.8–433.2) | 78.7 (78.4–79.1) | 46.9 (46.2–47.6) |
| xVDM, 18 hpf low perm. | 2 | 397.0 (396.9–397.1) | 73.4 (70.5–76.4) | 44.3 (42.1–46.5) |
| xVDM, 24 hpf | 2 | 396.5 (382.4–410.7) | 86.2 (85.9–86.5) | 50.2 (50.2–50.3) |
| VDM control, 24 hpf | 2 | 245.9 (244.7–247.2) | 28.6 (27.8–29.3) | 23.6 (23.1–24.1) |

| *(b) cDNA libraries* |  |  |  |  |
| --- | --- | --- | --- | --- |
| Condition | *n* | Accepted reads<sup>4</sup>(M) | Mol. hubs<sup>5</sup> (M) | Genes $\geq 10$ wt. support<sup>6</sup> |
| xVDM, 12 hpf | 2 | 329.2 (299.9–358.5) | 58.1 (57.4–58.8) | 23,572 (23,433–23,710) |
| xVDM, 18 hpf high perm. | 2 | 339.2 (317.6–360.8) | 71.2 (69.7–72.7) | 24,601 (24,457–24,745) |
| xVDM, 18 hpf low perm. | 2 | 337.4 (317.8–357.0) | 63.3 (63.2–63.4) | 24,080 (23,957–24,202) |
| xVDM, 24 hpf | 2 | 335.1 (307.8–362.3) | 77.6 (73.5–81.6) | 24,682 (24,577–24,787) |
| VDM control, 24 hpf | 2 | 100.7 (98.0–103.5) | 5.6 (5.5–5.8) | 16,934 (16,704–17,163) |

<sup>1</sup> Post-filter reads assigned to the UEI amplicon library.
<sup>2</sup> Distinct molecular nodes on the UEI library that participated in at least one retained linkage (“UEI-associated molecular nodes”); these are the primary UEI-side quantities reported here because the chemistry-side gain is expressed mainly as graph densification in the participating node set rather than as an increase in the number of distinct pairings alone.
<sup>3</sup> Subset of UEI-associated molecular nodes supported by $\geq 2$ reads.
<sup>4</sup> Post-filter reads assigned to the cDNA amplicon library.
<sup>5</sup> Molecular hubs with at least one assigned cDNA consensus; a library-yield count summing the 1-read, 2-read, and $\geq 3$ -read consensus bins. It is taken before gene-annotation filtering, before removal of genome-only and raw-sequence features, and before the row filter used to build the consolidated hub matrix.
<sup>6</sup> Genes whose accumulated weighted support across cDNA-bearing hubs reached at least 10.

**Table S3.** Read-depth-matched benchmark of xVDM versus transcript-anchored controls at 24 hpf. Values are medians across two embryos per chemistry and were obtained by rarefaction to the deepest common accepted-read depth shared by all four embryos for each amplicon library. The UEI library was compared at 204.8 million accepted reads and the cDNA library at 51.2 million accepted reads. Distinct pair counts are intentionally omitted as the primary UEI metric because the matched-depth chemistry gain is concentrated in the number of participating molecular nodes. Weighted gene support is fractional when a molecular hub carries more than one gene identity.

| Metric (24 hpf, matched depth) | VDM | xVDM | xVDM/VDM |
| --- | --- | --- | --- |
| *UEI amplicon library (204.8 million accepted reads)* |  |  |  |
| UEI-associated molecular nodes (M) | 27.8 | 69.0 | 2.5× |
| UEI-associated molecular nodes $\geq 2$ reads (M) | 22.6 | 38.1 | 1.7× |
| UEI-associated molecular nodes $\geq 3$ reads (M) | 19.7 | 27.7 | 1.4× |
| *cDNA amplicon library (51.2 million accepted reads)* |  |  |  |
| cDNA-bearing molecular hubs (M) | 5.3 | 32.6 | 6.2× |
| cDNA-bearing molecular hubs $\geq 2$ reads (M) | 4.6 | 11.3 | 2.5× |
| Genes $\geq 1$ weighted support | 23,428 | 26,210 | 1.1× |
| Genes $\geq 5$ weighted support | 18,562 | 23,312 | 1.3× |
| Genes $\geq 10$ weighted support | 15,738 | 21,592 | 1.4× |

### cDNA-hub co-detection analysis

We analyzed a consolidated cDNA matrix as a hub-level control, independent of inferred coordinates from the UEI-library embedding and independent of putative cell segmentation. This matrix is stricter than the cDNA hub yield reported in Table S2, which counts every molecular hub with at least one assigned cDNA consensus. The consolidated matrix was built from per-sample cDNA matrices after removing sub-consensuses with more than one retained gene assignment. During consolidation, a cDNA feature call was retained only if it had at least two supporting reads. A hub was kept only if at least two retained gene features remained. No protein-coding filter was applied in forming the matrix. The rRNA comparison below additionally required protein-coding signal so that rRNA-only hubs were excluded. After these filters, the matrix contained 52,280,187 cDNA hubs across eight embryos and 27,367 detected features, including 21,990 protein-coding features, 842 rRNA features, and 2 mitochondrial rRNA features. For the matched analyses, including the plotted panels and reported matched-hub counts, we drew a reproducible random subset of at most 250,000 cDNA hubs from each specimen. The biological specimen remained the unit of replication. The full consolidated matrix was used to report dataset size and feature content.

For the rRNA-containing hub comparison, matching was performed separately within each embryo. We first defined two mutually exclusive hub pools. The cytosolic-rRNA-containing pool contained hubs with at least one protein-coding feature and at least one cytosolic rRNA feature, but no mitochondrial rRNA feature. The mitochondrial-rRNA-containing pool contained hubs with at least one protein-coding feature and at least one mitochondrial rRNA feature, but no cytosolic rRNA feature. Hubs carrying both cytosolic and mitochondrial rRNA, or lacking protein-coding signal, were excluded from this comparison.

We then balanced the two pools for UMI type and hub complexity. Hub complexity was defined as the total number of detected features on the hub, with all hubs carrying six or more detected features placed into one capped bin. Within each embryo and within each UMI-type-by-complexity stratum, the larger pool was randomly downsampled to the size of the smaller pool. The retained strata were then concatenated across embryos. This produced 174,704 cytosolic-rRNA-containing hubs and 174,704 mitochondrial-rRNA-containing hubs for the gene-level comparison. For each protein-coding gene, presence or absence in the two balanced hub pools was compared within embryo, and embryo-level log odds ratios were combined by inverse-variance fixed-effect meta-analysis.

For each protein-coding gene, we converted gene presence in the two matched hub groups into a log odds ratio. Before computing the ratio, we added 0.5 to each cell of the 2 × 2 table. Let *a* and *b* denote the gene-present and gene-absent counts in the mitochondrial-rRNA group, and let *c* and *d* denote the corresponding counts in the cytosolic-rRNA group. The specimen-level log odds ratio was log[(*a*/*b*)/(*c*/*d*)], with sampling variance 1/*a* + 1/*b* + 1/*c* + 1/*d*. Specimen-level log odds ratios were combined by inverse-variance fixed-effect meta-analysis. *P* values are two-sided Wald *P* values from the fixed-effect *z* statistic, and *q* values are Benjamini–Hochberg adjusted over the tested protein-coding genes within each analysis. The plotted *q* values use the fixed-effect tests.

The fixed-effect analysis found 38 genes enriched in cytosolic-rRNA hubs and 19 genes enriched in mitochondrial-rRNA hubs at *q <* 0.05. Mitochondrial-rRNA hubs favored mitochondrial genes, including *mt-co1, mt-co3, mt-co2, mt-atp6, mt-nd6, mt-cyb, mt-nd2*, and *mt-nd5*. Cytosolic-rRNA hubs favored lipid and yolk genes, including *apoa1a, afp4, apoa1b, tfa, bhmt, apoeb*, and *fabp1b*.*1*. The per-embryo profiles and the 24 hpf minus 12 hpf comparison are shown in Fig. S6a,b. For Fig. S7, the same matched analysis was applied to the two 24 hpf embryos after replacing the rRNA compartment label with an anchor-gene label: hubs carrying *hbae3* or *ctslb* were compared with matched hubs lacking that anchor gene.

### DNA microscopy image inference

The likelihood model, the projected-gradient solver, and the three-range representative-pair sampler for the repulsive normalizer are inherited from ^14;15^. Three elements are reformulated: the local kernel that pre-conditions the spectral basis, the assembly of that basis from random rotations of rescaled eigenvectors, and the use of Infomap-derived aggregate cells as a post-reconstruction summary for slice-based registration.

Although the data record two molecular species (type-I and type-II UMIs), the inference treats the largest contiguous UEI matrix as an undirected weighted graph, in the same EASL-pruned form as in ^14^. Letting **N** denote the bipartite UEI count matrix and **C** = **N** + **N**^⊤^ its symmetrization, with 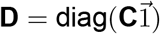, every major spectral and likelihood step below operates on **C** and its row-normalized form **D**^−1^**C**. The two-part labeling is preserved only for downstream gene annotation.

### Stabilized geodesic kernel

Let **Φ**^(1)^ collect the leading nonconstant right eigenvectors of **D**^−1^**C**, with corresponding eigenvalues *λ*_*k*_ ∈ [0, 1). As in ^14^, these modes are rescaled by inverse spectral gap,

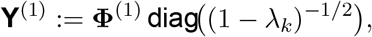

so that each retained dimension contributes equally to the quadratic part of the inference objective. This is the same “as-uniform-as-possible” criterion used to set rescaling factors in the earlier paper. The rows 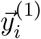 of **Y**^(1)^ are then stabilized by streamed random rotations followed by rank transformation. Many Haar-random rotations are drawn, each rotated coordinate is replaced by a centered rank score, and the leading directions of the accumulated collection are tracked by an incremental low-rank SVD without storing the full collection.

This is the same construction ^14^ used to inflate sub-sample-derived candidate solutions into an *E*_final_-dimensional spectral basis. Applied here to the full-data rescaled basis it serves the same purpose, namely to reduce the influence of extreme coordinates along any single spectral axis, and to spread that influence across many rotated views before the local neighborhoods used by the next step are formed. We continue to write the stabilized rows as 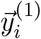 since the rotation-plus-rank operation is column-wise and preserves the row indexing of **Y**^(1)^.

Local neighborhoods are then formed in this stabilized space. For each node *i*, let **M**_*i*_ collect the displacement vectors 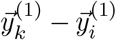 for *k* in the *k*-nearest neighbors of *i*. For each observed edge (*i, j*) with weight *N*_*ij*_, the edge displacement 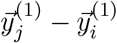 is fit by a nonnegative local mixture,

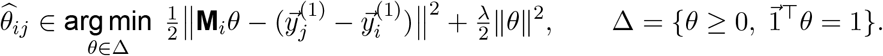

The simplex constraint keeps the fit interpolative (each edge is expressed as a convex combination of locally observable directions), and the ridge term damps the weights when nearby directions are nearly collinear. The row of the smoothing matrix for node *i* is then 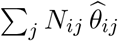, placed on the columns indexed by *N* (*i*). Symmetrization and row normalization give the smoothing matrix **W**.

This object plays exactly the role of the geodesic Gaussian kernel 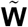 in ^14^. There, neighborhood geometry was summarized by tangent-corrected pairwise distances passed through a Gaussian RBF. Here it is captured by a constrained least-squares fit on raw displacement vectors. The two share sparsity pattern, role (left/right smoothing operator for the next spectral problem), and motivation, namely to encode locally reliable graph geometry in a kernel that is stable under the noise and density variations seen in large UEI graphs.

### Transport-smoothed spectral pass

The smoothing matrix **W** is applied on both sides of the symmetrized graph. At the operator level this corresponds to the product

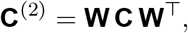

though in practice **C**^(2)^ is never assembled: a block-Krylov solver applies **W** and **W**^⊤^ as left/right operators on **C**, with the appropriate row normalization absorbed into a single diagonal rescaling. The leading nonconstant modes of the row-normalized form of **C**^(2)^ are computed in this way, rescaled again by their inverse spectral gap, and passed through the same rotation-plus-rank stabilization used in the first pass.

The rank-stabilized columns are then interleaved, in groups matching the embedding dimension, with the corresponding smoothed eigenvectors before a column-wise modified Gram–Schmidt produces the final orthonormal basis **Q**. Interleaving in groups of the embedding dimension ensures that neither view dominates the leading directions of **Q**. The rank view damps sensitivity to local density and outlying coordinates, while the raw smoothed view preserves large-scale structure that ranks alone may flatten.

The procedure differs from ^14^ in how the local kernel is learned and in how candidate directions are produced for the basis, but not in the overall numerical strategy. Both versions fold a locally adaptive smoother into the spectral problem and then stabilize the resulting candidate basis by random rotations and ranks. The earlier paper rotated a stack of sub-sample interpolations into the basis, whereas the present version rotates the rescaled full-data basis. The two are interchangeable in slot, and on the data sizes considered here the full-data route is preferable because it removes the interpolation step entirely.

### Reconstruction by likelihood maximization

Coordinates are written as **X** = **QB**, with **Q** the orthonormal basis from the diffusion-transport-smoothed spectral pass and **B** the active coefficients. The solver is the incremental projected gradient descent of ^14^: basis directions enter in increasing spectral order, each newly activated block receives a local second-order trust update, and is then refined jointly with the already-active block. As in that paper the interaction-decay law is held fixed during this refinement.

Let 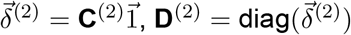, and 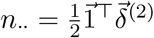. The reduced-space objective is

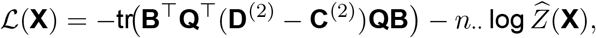

the same multinomial log-likelihood as in ^14^, now evaluated on the diffusion-transport-smoothed graph **C**^(2)^ and projected into the smoothed spectral basis **Q**.

The repulsive normalizer 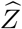 uses the same three-range representative-pair sampler. Sampled pairs are refreshed between solves so each inner solve differentiates a fixed sampled objective, matching the procedure in ^14^. The only modification is that endpoint weights in the sampled normalizer are derived from diffusion-transport-smoothed connectivity,

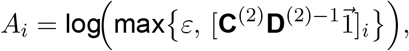

so that nodes that remain strongly connected after smoothing carry proportionally greater weight, in the same form as the earlier paper’s *A*_*i*_ ← log ∑_*j*_ *n*_*ij*_/*n*_·*j*_ but with **C**^(2)^ replacing **N**.

Gradients and Hessian–vector products are taken with respect to **B**, with ∇_**B**_ = **Q**^⊤^ ∇_**x**._ and 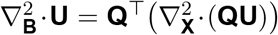. The attractive term contributes the projected graph Laplacian **Q**^⊤^ (**D**^(2)^ − **C**^(2)^)**Q**, and the repulsive term contributes representative-pair sums.

Note that ℒ depends only on pairwise differences 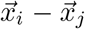, so reconstructions are determined up to a global rigid motion. We mean-center reported coordinates and align by Procrustes when comparing across runs or to ground truth.

### Voronoi compartment simulation

The accuracy curves and scatter plots in Fig. 1 are based on synthetic graphs with known compartment labels. Molecules are sampled uniformly inside a three-dimensional Voronoi tessellation, inheriting a ground-truth identity from the region containing them.

Compartment structure is imposed through a spatially varying effective diffusivity *κ*(**x**). A baseline intracellular value is modulated by quasi-periodic spatial modes and per-cell stochastic offsets, and near each Voronoi face log *κ* is decremented by a Gaussian membrane barrier *β*_mem_ exp(−(*δ*/*w*_mem_)^2^), with *δ* the distance to the nearest face. The path-averaged value 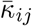 along a candidate pair (*i, j*) then sets the screened interaction length 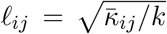, with *k* a uniform degradation rate. Pairs that cross a membrane have a shorter *ℓ*_*ij*_ and are less likely to link.

Short-range associations follow a Gaussian overlap kernel set by the nanoball radius. Long-range associations follow the screened Yukawa kernel ∝ *r*^−1^ exp( − *r*/*ℓ*_*ij*_). Conditional on a fixed total yield, counts over candidate pairs are multinomial in the kernel weights, with source counts drawn from row marginals and targets sampled within bounded neighborhoods.

#### Connection-spread-function analysis

For each reconstruction we computed the UEI-weighted histogram of inferred edge distances in GSE coordinates,

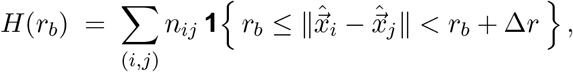

where *n*_*ij*_ is the retained UEI count between molecular nodes *i* and *j* and 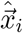 is the inferred coordinate of node *i*. Normalized to unit area over the plotted range, this histogram is the connection-spread function (CSF). Distances are in diffusion units of the GSE reconstruction.

Under locally uniform node density and Gaussian molecular dispersion in three dimensions, the radial CSF follows a Maxwell–Boltzmann form ^14^,

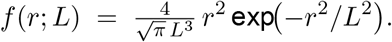

For plotting and fitting, the histogram was restricted to the displayed range *r <* 3 diffusion units and normalized to unit area over that range. Model curves were renormalized over the same range before fitting. The fraction of total UEI mass at *r* ≥ 3 diffusion units was not included in the normalization and is reported separately in Fig. 3h.

The CSF captures how rapidly retained molecular proximity information decays in the inferred geometry, providing a connectivity-based resolution proxy. The per-node localization scale predicted by the DNA-microscopy likelihood approximation of ^14^ is 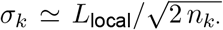 with *n*_*k*·_ = ∑_*j*_ *n*_*kj*_, conditional on the inferred positions of UEI-connected neighbors. We use this expression only as an interpretive bridge to the prior VDM framework, because the native xVDM observable is a molecular proximity graph rather than a set of independent point positions.

### Segmentation: simulation benchmark and xVDM cell calling

For the Fig. 1 benchmark, performance is normalized mutual information against the true Voronoi labels. Nodes a method leaves unlabeled are excluded from its score. The UMAP baseline is graph-based: from the symmetrized UEI graph we take the 15 leading eigenvectors of the symmetric normalized adjacency matrix (dropping the trivial one), standardize them, and run UMAP in three dimensions with 30 neighbors and cosine distance, with `min_dist`=0.1 and 0.99 (Fig. 1k,m). Two comparisons follow. In the embedding-controlled comparison the same segmenter (HDBSCAN) is applied after each embedding, so score differences reflect the embedding alone, using the `min_dist`=0.99 UMAP view. Here GSE outperforms UMAP, which broadens compartment density. Separately, GSE is scored by its connectivity route, Infomap on the diffusion-transport graph, so panels n–q show GSE under both HDBSCAN and Infomap. In all cases unlabeled nodes are reassigned only when a strict majority of their symmetrized-graph neighbors share a label. The node-density panels (n,o) jointly vary total node count and cell number. The barrier panels (p,q) vary cell number and the membrane-to-intracellular diffusivity ratio. The decoherence radius (r) is the spatial spread at which the mean Procrustes RMSD between matched local neighborhoods exceeds 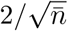, where 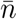 is the mean symmetrized UEI count per node.

For xVDM cell calling, the primary labels were obtained by running Infomap on a sparse diffusion-transport graph derived from the final GSE reconstruction. Nodes initially left unlabeled were reassigned only when a strict majority of their graph neighbors shared a label. The resulting labels were split by connected components in the raw symmetrized UEI graph, and components below the minimum size threshold were demoted to unlabeled. Genetic annotation was aggregated after these connectivity-derived labels were assigned.

### Reference-guided registration

The xVDM reconstruction gives a three-dimensional map in diffusion units, but its orientation is arbitrary: the reconstructed coordinates are fixed only up to rotation, reflection, and translation (*SI: Reconstruction by likelihood maximization*). To align this map to a two-dimensional tissue reference, we used shared gene expression as a coordinate system both datasets can read. We assigned each xVDM aggregate cell to a site in a matched-stage Stereo-seq slice. The assigned slice coordinates are used for comparison, display, and held-out marker tests. They are not fed back into the molecule-level embedding.

In the main inference pipeline, aggregate cells were defined after the final embedding and graph-based cell call, and the pipeline then ran one fixed registration setting. In the held-out marker benchmark, the same aggregate cells were registered repeatedly as the marker budget was varied. In both cases the reference slice supplies positions 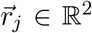 and gene counts 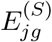 for slice sites *j* = 1, …, *n*_*S*_, and the reconstruction supplies gene counts 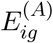 for aggregate cells *i* = 1, …, *n*_*A*_, summed from each cell’s molecular hubs. Registration returns a slice site for each aggregate cell,

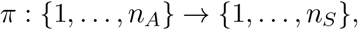

and the associated slice coordinates

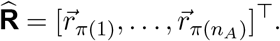

#### Gene-pole fields

A gene-pole field is one expression axis that compares a small gene group *A*_*p*_ with an opposing group *B*_*p*_. The groups are chosen from the matched-stage slice so that the two sides mark distinct, internally coherent regions and the resulting *P* axes are weakly correlated across the slice. Genes are not reused across axes in the held-out benchmark. For pair *p* at slice site *j*, we sum expression on each side and smooth those sums over nearby slice sites,

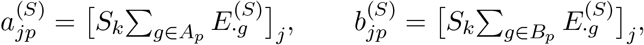

and form the ratio

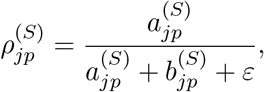

with neutral value 1/2 where both sides vanish. The same calculation on xVDM aggregate cells, using aggregate-cell cDNA counts smoothed in the reconstructed embedding, gives 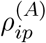. Each field is rank-transformed to [0, 1] within its own assay, so the comparison depends on spatial ordering rather than absolute expression scale.

#### Capacity-constrained slice assignment

Each slice site *j* is given an integer capacity *c*_*j*_. In the configuration used here, the capacity is proportional to the total slice expression mass

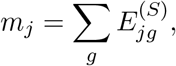

normalized so that ∑_*j*_ *c*_*j*_ = *n*_*A*_. Slice sites with more total RNA therefore receive more aggregate cells. Let 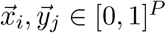 be the gene-pole feature vectors for aggregate cell *i* and slice site *j*. The cost of assigning *i* to *j* is a Huber sum over fields,

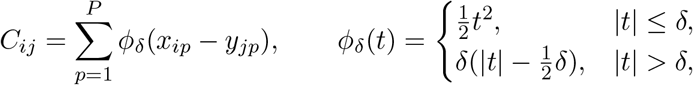

whose quadratic core preserves sensitivity where the assays agree and whose linear tail keeps a single under-detected pole from vetoing an otherwise sensible placement. The base map is the minimum-cost assignment under the capacities,

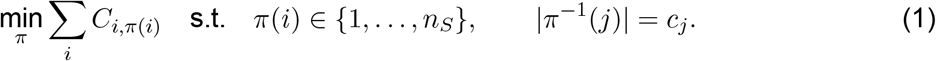

For speed, each aggregate cell is first restricted to a small set of nearby slice sites in gene-pole feature space. This candidate set is grown until a capacity-respecting assignment exists, and the base map is the exact optimum on that sparse candidate graph.

#### Graph refinement

The base map scores aggregate cells one at a time, coupled only through the global capacity constraint. The refinement step asks aggregate cells that are close in the xVDM connectivity graph to land near one another on the slice, adding the penalty

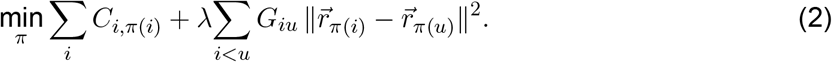

Here *G* is the aggregate-cell graph obtained from the diffusion-transport graph by summing weights within and between aggregate cells, then symmetrizing and normalizing. The refinement therefore reuses the connectivity learned during reconstruction, now at the aggregate-cell scale. The penalty is linearized around the current assignment, and the assignment is solved again under the same capacities. The reported runs used two refinement rounds with 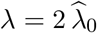, where 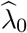 is an internally estimated weight that places the feature and graph costs on comparable scales. The factor of two is the only exposed setting, and a factor of zero recovers the feature-only map. Because the penalty depends only on distances among assigned slice coordinates, it encourages local consistency without imposing a separate rotation or reflection.

#### Held-out marker benchmark

For each dataset, a library of gene-pole pairs was built from the matched-stage Stereo-seq slice, with up to five ranked genes per side. For a budget of *B* genes per side and *g* genes per pole, the benchmark used *B*/*g* training pairs when *B* was divisible by *g*. The runs reported here used *B* ∈ { 5, 10 } and *g* ∈ { 1, 5 }, giving four settings: five single-gene pairs, one five-gene pair, ten single-gene pairs, and two five-gene pairs. Three folds were run with three held-out pairs per fold, and held-out scoring used single-gene pairs whose genes were disjoint from the fitted markers. Within each run, training and held-out fields were projected onto a principal-component basis defined only by the target slice fields. The PC1 correlograms compare target and predicted scores for the base and refined maps. A held-out point near the diagonal means registration transferred spatial structure beyond the fields used for fitting.

#### Registration output

Registration returns the slice site assigned to each aggregate cell and the corresponding raw slice coordinates for the base and refined maps. These outputs support direct coordinate plots, the Fig. 3 benchmark panels, and the Fig. 4a mapping views. They do not replace the molecule-level xVDM coordinates from the reconstruction.

#### Gene-level compactness and external spatial autocorrelation

For Fig. 4c–e, we compared a gene-level spatial statistic from xVDM with an independently measured spatial statistic from the matched-stage Stereo-seq reference. For each stage, candidate genes were required to be present in all selected stage-matched xVDM specimens and to pass a minimum external Stereo-seq detection fraction of 0.02. If more than 3,000 genes passed these filters, genes were selected by stratified sampling across the external Moran’s-*I* ordering, taking high-autocorrelation, low-autocorrelation, and intermediate genes for balanced coverage of the rank range.

External spatial autocorrelation was measured as Moran’s *I* in the matched-stage Stereo-seq slice, using a six-nearest-neighbor graph over the slice’s physical two-dimensional coordinates. Independently, for each selected gene and each stage-matched xVDM specimen, high-expressing aggregate cells were defined by the upper expression quantile, using the 80th percentile of log-normalized expression. For sparse genes whose 80th percentile was zero, all detected cells were used as the high-expression set. A specimen-level compactness score was computed only when the gene was detected in at least 2% of aggregate cells and the high-expression set contained at least 25 cells.

Gene compactness was defined as

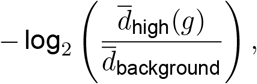

where 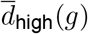 is the mean pairwise GSE distance among high-expressing aggregate cells for gene *g*, and 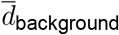 is the mean pairwise distance among aggregate cells in the same specimen. Larger values therefore indicate that high-expressing cells for that gene are more compact than the specimen-wide background. Finite specimen-level compactness scores were averaged across the stage-matched xVDM specimens.

Before correlation analysis, external Moran’s *I* and xVDM compactness were each residualized against six covariates: external mean expression, external detection frequency, external expression variance, xVDM mean expression, xVDM detection frequency, and xVDM expression variance. Residuals were transformed to rank-normal scores for visualization, and Spearman correlations were computed between the residualized external Moran’s-*I* values and residualized xVDM compactness values.

### Protein target hub and neighborhood analysis in human tonsil

Protein target UEI and cDNA libraries were generated as described (*SI: Human tonsil tissue preparation and antibody-oligonucleotide staining*), sequenced, and analyzed in a manner equivalent to other xVDM data. Antibody-derived inserts were parsed for recognized CD3, CD9, IgD, and CD24 epitope motifs, after canonicalizing motif aliases. Only inserts containing exactly two recognized epitope motifs were used for co-localization analysis. Each retained insert was assigned one unordered epitope-pair label, such as CD3/CD3, CD3/IgD, or IgD/IgD.

Hub-level tests compared hubs within the same exact number *d* of retained epitope-pair inserts. Within each exact-*d* stratum, the permutation null shuffled epitope-pair labels over the fixed insert slots, preserving the number of hubs, the number of retained inserts per hub, and the overall abundance of each epitope-pair label. *P* values were empirical label-shuffle tail probabilities, and Benjamini–Hochberg *q* values were computed within the corresponding exact-*d* stratum. For Fig. S10a,d, a heat-map box was counted only when the compared epitopes could be assigned to two distinct retained inserts on the same hub. For Fig. S10b,e, the analysis was restricted to homo-epitope insert labels such as CD3/CD3 and IgD/IgD. For Fig. S10c–h, neighborhood analyses were performed within the exact one-retained-insert stratum; Infomap-cluster maps were colored by *K* = 30 nearest-neighbor-smoothed log_2_ relative enrichment of CD3/CD3 versus IgD/IgD.

